# Clonal *Parabacteroides* from Gut Microfistulous Tracts as Transmissible Cytotoxic Succinate-Commensal Model of Crohn’s Disease Complications

**DOI:** 10.1101/2024.01.09.574896

**Authors:** Vaidhvi Singh, Gail West, Claudio Fiocchi, Caryn E. Good, Jeffry Katz, Michael R Jacobs, Armand Earl Ko Dichosa, Chris Flask, Mathew Wesolowski, Cassidy McColl, Brandon Grubb, Sadia Ahmed, Nicholas C. Bank, Kavitha Thamma, Ilya Bederman, Bernadette Erokwu, Xinlin Yang, Mark S. Sundrud, Paola Menghini, Abigail Raffner Basson, Jessica Ezeji, Satish E Viswanath, Alida Veloo, David B. Sykes, Fabio Cominelli, Alex Rodriguez-Palacios

## Abstract

Crohn’s disease (CD) has been traditionally viewed as a chronic inflammatory disease that cause gut wall thickening and complications, including fistulas, by mechanisms not understood. By focusing on *Parabacteroides distasonis* (presumed modern succinate-producing commensal probiotic), recovered from intestinal microfistulous tracts (cavernous fistulous micropathologies CavFT proposed as intermediate between ‘mucosal fissures’ and ‘fistulas’) in two patients that required surgery to remove CD-damaged ilea, we demonstrate that such isolates exert pathogenic/pathobiont roles in mouse models of CD. Our isolates are clonally-related; potentially emerging as transmissible in the community and mice; proinflammatory and adapted to the ileum of germ-free mice prone to CD-like ileitis (SAMP1/YitFc) but not healthy mice (C57BL/6J), and cytotoxic/ATP-depleting to HoxB8-immortalized bone marrow derived myeloid cells from SAMP1/YitFc mice when concurrently exposed to succinate and extracts from CavFT-derived *E. coli*, but not to cells from healthy mice. With unique genomic features supporting recent genetic exchange with *Bacteroides fragilis*-BGF539, evidence of international presence in primarily human metagenome databases, these CavFT *Pdis* isolates could represent to a new opportunistic *Parabacteroides* species, or subspecies (‘*cavitamuralis’*) adapted to microfistulous niches in CD.

---

Crohn’s Disease (**CD**) is a chronic inflammatory disease of the bowel with no known cure^1^ that affects millions of people worldwide. Unfortunately, as disease progresses, most patients require surgery to remove thickened and affected bowel and to treat complications, such as perforations and fistula formation, which is known as fistulizing CD. In context, fistulas are macroscopic cavitating inflammatory tracts that interconnect organs, but what precedes their formation is unknown.

Traditionally, pathologist have recognized the presence of microscopic lesions that initiate in the gut mucosa in CD and manifest as fissures that resemble small tears (or ‘cracks in a rock’) that start in the mucosal lining of the intestine and that penetrate perpendicularly a few micrometers into the submucosa^1,2^. Their relevance in disease is unknown, but recent studies we conducted in surgical specimens revealed their presence differs from that other unrecognized more structured cavitating microlesions we discovered in CD. By studying the three-dimensional microscopic architecture of the gut-wall in resected ileal tissue from patients undergoing intestinal surgery, we recently discovered that such cavitating microfistulous structures resemble geological cavern formation^2^, in which submucosal ‘sinkhole’ liquefactive lesions lead to **Cav**ernous **F**istulous micro**T**racts (**CavFT**) that are non-discernable by the naked eye or histology^2^. Through DNA and culturomics, we have identified numerous bacterial species that frequently inhabit CavFT, particularly *Bacteroidota,* including *Parabacteroides distasonis (Pdis)* (**NCBI BioProject**: PRJNA224116)^3,4^, which collectively indicate that bacteria could play a perpetuating inflammatory or modulating role in such intramural microniches.

Crohn’s disease was first described at the beginning of the last century, as regional ileitis by Crohn, Ginzburg and Oppenheimer in a case-series presented at a medical meeting in 1932^5, 6^. The main objective of this case-series and experimental study is to report the complete genome analysis of two *Pdis* isolates from intestinal microfistulous tracts in CD patients, in an international epidemiological context, to differentiate cross-contamination from clonality and potential person-person transmissibility (in hospital vs. community), and to determine experimentally if these commensal bacteria could induce CD-like ileitis in healthy mice or worsen the severity of ileitis in mice prone to CD-ileitis.

The species *P. distasonis* (including reference strain ATCC8503)^7^ has been purported as a potential probiotic species for anticancer and anti-obesity functions in recent years^8–11^. However, both clinical studies in serious medical infections and experimental studies in animals have shown that *P. distasonis* species are pathogenic, or an opportunistic pathobiont in humans, with and without inflammatory bowel disease (IBD)^12–14^.

Herein, genomic analysis revealed the two *Pdis* isolates in this CD report are clonal and have unique genomic features of exchange with *Bacteroides fragilis*, indicating this is novel lineage potentially adapted to CD micro-niches^15^ and suggestive of transmission in the community. Experiments on CD-prone (SAMP1/YitFc) ^16^ and healthy mice (C57BL/6J) showed these *Pdis* worsen ileitis in susceptible mice but not in healthy mice. Interestingly, *Pdis*, known for its anti-inflammatory properties and lacking LPS due to O-antigen operon fragmentation^15^, was cytotoxic to immortalized bone marrow derived progenitor myeloid cells from CD-prone mice, especially if combined with extracts from *E. coli* also derived from CavFTs.

## Results

### CavFT lesions as intermediate liking model between fissures and fistulas in Crohn’s Disease

To contextualize the pathological structure of CavFT lesions in humans, and further expand our knowledge derived from the study of these gut wall lesions using stereomicroscopy^2^, we developed a high-resolution 100-µm-voxel *ex-vivo* MRI imaging technique to illustrate how inflammation linked to CavFT transects all the gut wall layers reaching the serosal/peritoneal layers. A few features distinguish CavFTs from fissures and fistulas. Conceptually, as shown in **Figure 1A**, CavFTs originate from the mucosa lining as fissures, or as liquefactive lesions forming under the epithelium and extend or penetrate deep into the bowel within muscle layers and extend transversally along the intestinal axis within muscle bundles creating complex microlesions that create perforations through the gut wall in inflamed areas, running often underneath healthy mucosa creating fistulous microtracts that reach the serosa, peritoneal cavity and mesenteric fat (**Figures 1A-F & Supplementary Figure 1**). This understudied mechanism for bacteria translocation^17^ could help explain why recent culturomic studies in obese individuals have yielded the isolation of commensal *Pdis* from human mesenteric fat in the peritoneal cavity, and the identification of succinate (a Krebs’ cycle byproduct) as a mitigating factor in obesity^18^. Because CavFT often connects the gut lumen with the serosa/peritoneal cavity and contains live bacteria, we earlier proposed that CavFT-bacteria could promote inflammation and extend cavitations and in some cases be the precedent to fistula formation^2^.

**Figure 1.**
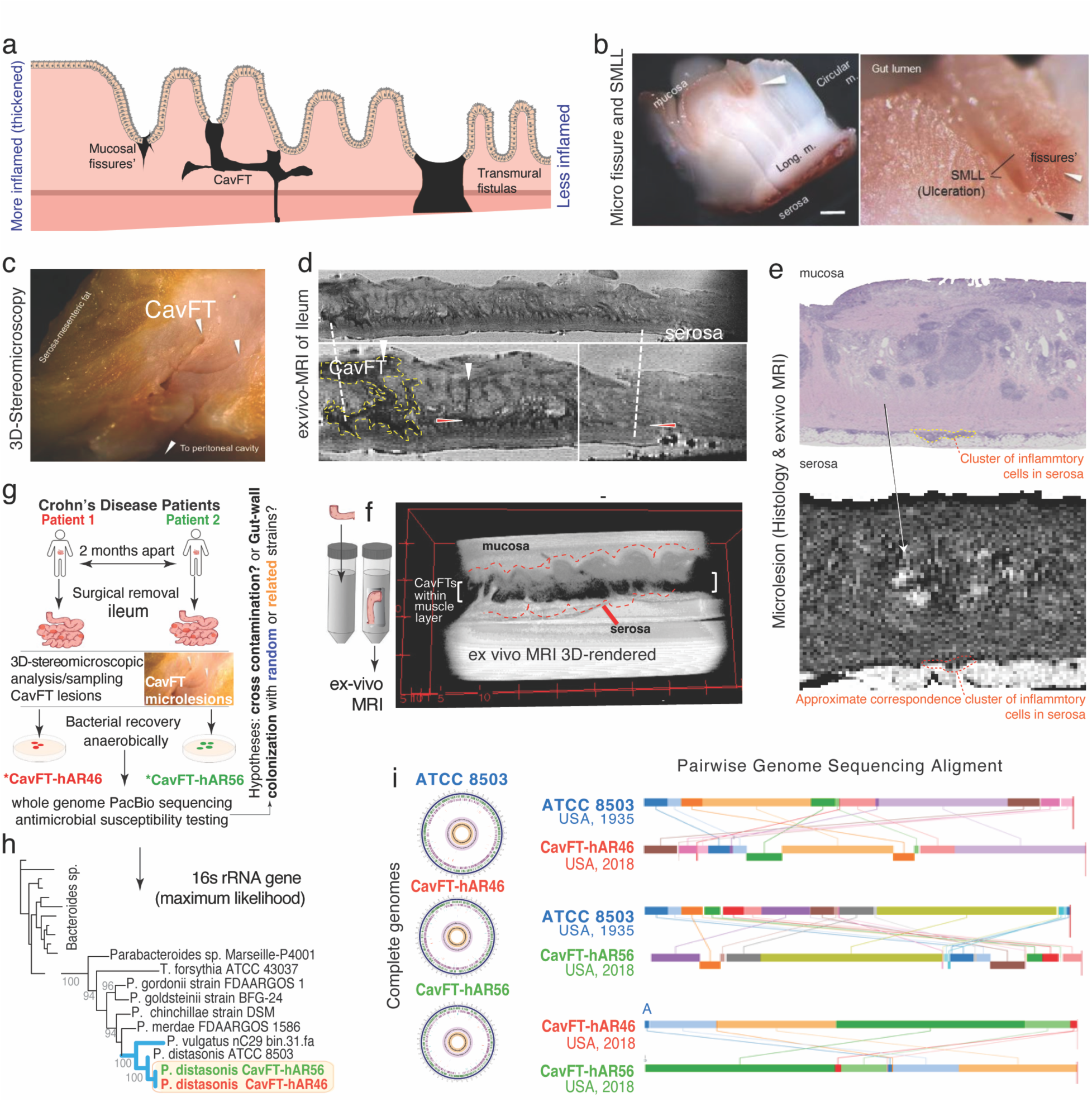
Structural overview of cavitating microfistulous tracts (CavFT) and genomics of *P. distasonis* from CavFT in CD patients. **a)** Graphical representation of differences between mucosal fissure, CavFT and transmural fistulas. Hierarchically, ‘sinkhole’ liquefactive lesions on the mucosa often lead to penetrating micro-fissures and ‘cavernous fistulous tract (CavFT/*cafta*/) microlesions’. **b)** 3D-Stereomicroscopic (SM) image illustrating a liquefactive lesion (SMLL) indicative of epithelium ulceration and a micro fissure in a colon sample from patient with CD fixed in formalin^2^. See **Supplementary Table 2** for detail definitions of all cavitating stages found in IBD^2^. **c)** 3D-SM and **d)** axial magnetic resonance imaging (MRI) of inflamed ileum with regional thickening and intramural hypoechoic/dense lesions (left, thicker). **e**) Close up of MRI-detectable intramural lesions (granulomatous) and approximate corresponding H&E staining of lesions, illustrating agreement and serosal hyper echogenicity (cluster of inflammatory cells). **f**) 3D-rendered profile of stack 100um voxel MRI images taken 1mm apart of ileum of another patient (imaged immersed in hyperechoic solution to illustrate contrast). To visualize the penetrating inflammatory tracts (horizontal & vertical lesions) the brightness/contrast have been adjusted to decreased pixel intensity in submucosa. Notice fistulous/penetrating CavFT-like lesions connecting the mucosa and peritoneum/serosa (resembling icicles) within the transparent submucosa. **g**) Overview of case-series with culture and identification of CavFT *P distasonis*. Inset photo, CavFT reaches the peritoneum^2^. **h)** 16s RNA gene phylogenetics. 1000 bootstrap value. CavFT isolated cluster together. **i)** Circle plots of the complete *Pdis* genomes in this study (publicly available at BV-BRC database). Line plots, Mauve genome pairwise alignment of *Pdis* ATCC 8503 (1935), CavFT-hAR46 and -hAR56 (2017). Distinct colors illustrate rearranged genome fragments. CavFT-pairwise shows few rearrangements, indicating strains have evolved separately.

### Genomes of P. distasonis from CavFT/Crohn’s disease patients imply community transmission

To elucidate the role of *Bacteroidota* in CD-CavFT, herein, we focused on characterizing and analyzing the genomic epidemiology of *Pdis* from two CavFTs in unrelated CD patients. These patients underwent surgical removal of their affected ileum two months apart (*Patient1/*CavFT-hAR46 & Patient2/CavFT-hAR56, see 16S/23S ribosomal sequence analysis in **Figures 1G-H & Supplementary Figure 2A-B**).

Although the likelihood of isolating rarely cultured bacteria (*Pdis)* from two consecutive patients is very low (1 in 59000, p<0.0001), we next determined whether these isolates resulted from cross-contamination in the hospital or laboratory, or if they were different and clonally-related. For this purpose, we use PacBio long-range sequencing to obtain the complete genome of Patient2/CavFT-hAR56, the complete re-sequenced genome of Patient1/CavFT-hAR46 and compared these two with the original PacBio complete genome report^4^ of Patient1/CavFT-hAR46. Genome analysis revealed the CavFT isolates were nearly identical among themselves (99.97%), yet more different from other *Pdis* strains (77.9-84.9%), ruling out cross-contamination, and supporting the notion of clonality with potential patient-to-patient transmission in the community and not the hospital where surgeries were performed or the laboratories where the surgical specimens were processed (**Supplementary Table 1**). Based on single nucleotide polymorphisms (SNP-mutations), CavFT isolates differed by >9 SNPs (1314 Gap-bases, 86 indels, Edge Bioinformatics, Los Alamos National Laboratory), which is greater than the 3-SNP-threshold used to define hospital transmissions (four months apart^19–21^) for *Clostridioides difficile* (**Supplementary Figure 3**), ruling out recent hospital person-person transmission. Compared to the reference *Pdis* strain ATCC-8503, which was isolated from human feces in 1935, extended protein sequence analysis with 100 proteins illustrates the strains are different, yet together form a unique clone that seems evolutionary distant from other clinical fully sequenced isolates (**Supplementary Figure 2C**).

### Genome rearrangement analysis shows P. distasonis are ‘bombarded’ with Bacteroides DNA

To further delineate the clonality of these two CavFT-*Pdis* isolates, we looked at the structural profiles of the complete bacterial genomes. Since single-nucleotide polymorphisms (SNP) are overly sensitive to make inferences about evolution, we next conducted whole-genome pairwise analysis to assess ‘gnome rearrangements’, which are less frequent and more conserved than SNP mutations^22–24^.

Of interest, our *Pdis* CavFT isolates share many structural genomic features and differ only in few DNA rearrangements, compared to the genome arrangement profile of our strains matched to the *Pdis* reference strain ATCC8503, demonstrating that these two CavFT *Pdis* have evolved in parallel, but still belong to a unique cluster in the context of all other available *Pdis* genomes (**Figure 1I, Supplementary Figure 4**).

Then, we quantified the DNA similarity of the *Pdis* genomes to that of other representative bacterial species from other *Bacteroidota* genera that we have isolated from CavFTs^4, 25^. We achieved this by conducting genome pairwise alignments and by listing the coordinates and DNA sizes of all the DNA sequences (inserts) that were similar (shared) between our *Pdis* and the complete genomes of other bacteria of interest identified in our culture analysis or by NCBI blast in this study. By examining the DNA sequences of the rearranged genome fragments, we found that a vast proportion of the rearranged CavFT sequences (31-33.9%, in fragments of <24000bp) matched various *Bacteroides* species, being statistically more common for *Bacteroides* in number and sizes compared to other genera within the phylum *Bacteroidota*, and over other *Pdis strains* (ANOVA p<0.001).

The distribution of size and number of such sequence fragments share between *Pdis* and the comparator species is illustrated as a ‘claw plot’ in **Figure 2A**. These shared DNA statistical analyses led us to hypothesize that *Pdis* has favored the exchange of DNA with *Bacteroides* over other genera throughout evolution (*e.g., Alistipes*, which appear less frequently and at smaller sizes in *Pdis*, **Figure 2A**).

**Figure 2.**
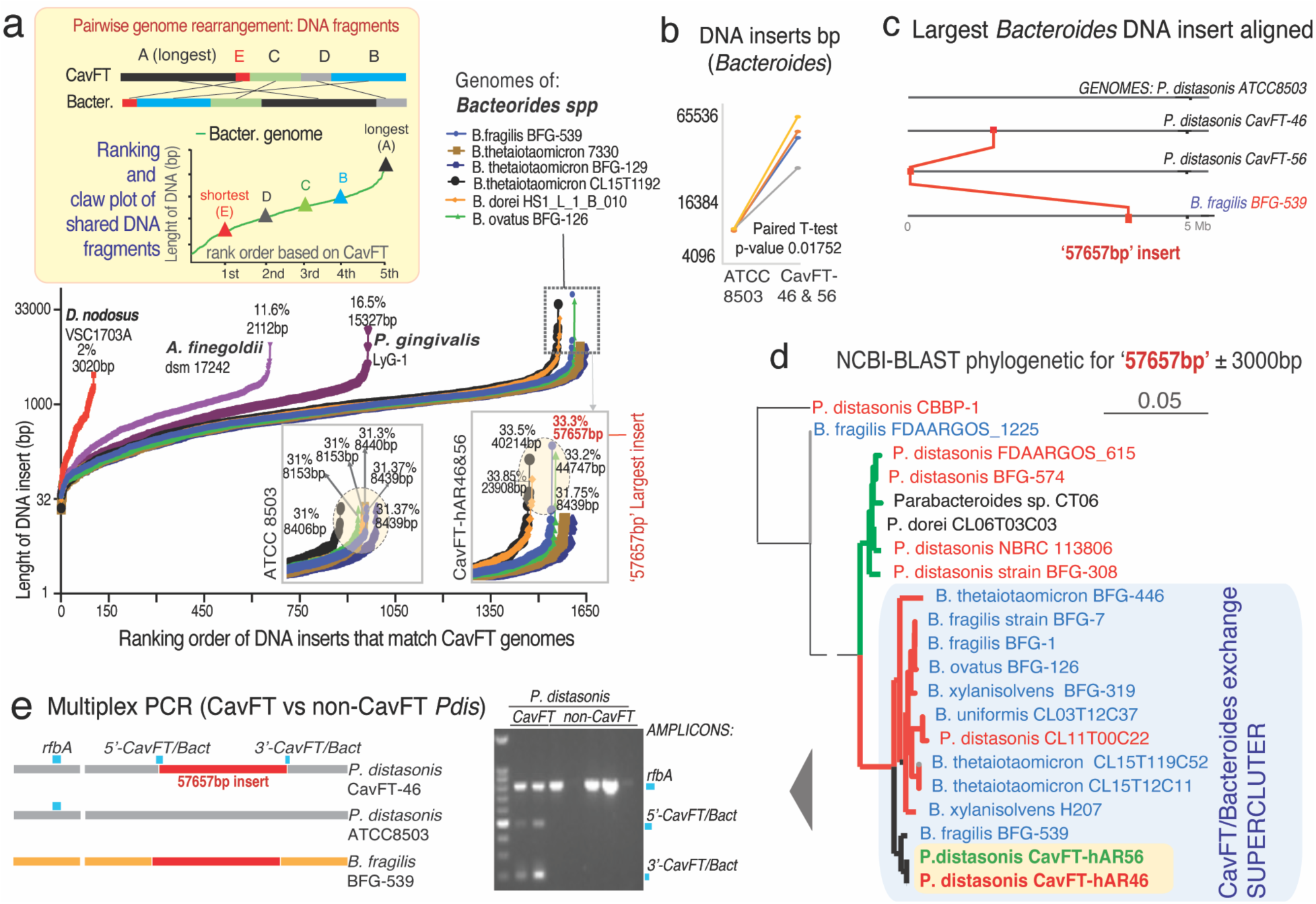
*Parabacteroides distasonis* CavFT strains are not cross-contamination but clonal members of a potential new subspecies (*cavitamuralis*). **a)** Inset, overview of experiment. ‘Claw plot’ illustrating the ranking of DNA fragments that CavFT-hAR46 shares with other genomes. *Bacteroides* spp. contribute to the largest number/sizes of foreign DNA in CavFT-hAR46**. b)** Line plot inset, CavFT has 3-7X larger *Bacteroides* inserts than ATCC-8503 (**Supplementary Figure 5**). **c)** The largest foreign DNA insert in CavFTs is 57657bp from *B. fragilis* BFG-539, shown aligned with other genomes. **d)** NCBI-Blast Phylogenetic tree of this CavFT/*Bacteroides* insert (NCBI: NZ_CP124739.1: Region 15183-72840)±3,000bp reveals CavFT strains are novel, unique and distinguishable in the NCBI database. Emerging ‘CavFT/*Bacteroides* exchange supercluster’ proposed. **e)** Multiplex PCR primers to study the epidemiology of ‘CavFT/*Bacteroides* exchange cluster’. Schematics and gel show expected amplicons for CavFT-*Bacteroides* insertions 5’-CavFT/bact (3’-CavFT/bact and *rfbA-typing*).

In perspective, such DNA sharing (or presumable exchange) resembles a pattern of ‘systematic bombardment of *Bacteroides* DNA’ into *Pdis* genomes, which we recently showed is not random. We also proposed in other *Pdis* that such bombardment could be a mechanism by which numerous antigenic operons are fragmented and rendered non-functional in *Bacteroidota*^15, 26–28^, including impairing LPS in *Pdis* CavFT, and enabling a silent form of ‘invasion’ of gut wall by those commensal bacteria^26^.

### The largest foreign DNA fragments in CavFT P. distasonis belong to Bacteroides fragilis

Since we have also isolated numerous *Bacteroides* from CavFT, including *B. fragilis, B. uniformis, B. ovatus, B. thetaiotaomicron*, and *Phocaeicola dorei*^2^, we propose that micro-niches such as CavFT could be favoring the genomic exchange among CavFT co-habiting species promoting niche-specific bacterial adaptation, evolution and persistence in patients, in a fashion that could resemble how species adapt to locally unique environments (*e.g.* mountain caverns).

Supporting such a possibility, we next searched the most distinctive *Pdis*/*Bacteroides* pairwise DNA-shared fragment profiles for each pairwise genome rearrangement analysis and discovered that the largest *Bacteroides* DNA insertions in *Pdis* (e.g., 57,657bp, >99.9%) is a DNA fragment that is identical to *B. fragilis* BFG539 (NCBI: CP103203.1) were only identified in CavFT *Pdis* and not other *Pdis* strains. Statistically, the top 10 largest *Bacteroides* insertions in CavFT strains were also 7-fold larger than the largest insertions in the *Pdis* reference strain ATCC8503; T-test P=0.017; **Figure 2B-C)**. Interestingly, such sharing of large DNA fragments seen between CavFT and *B. fragilis*, also occurs among other *Pdis* and *Bacteroides*, for other fragment sizes (∼30,000bp, *B. ovatus* BFG-139) which can be visualized as a separate phylogenetic clade (*Pdis/Bacteroidota* supercluster, **Figure 2D**). To further verify that such sequences in our *Pdis* genomes encode known functions in *Bacteroides* (and not *Pdis*), we fragmented *in silico* the largest DNA insertions into 250bp and used BV-BRC metagenomic pipelines to annotate such a collection of 250bp fragments. This functional metagenomic analysis confirmed that 71% of the DNA sequences perfectly aligned between *Pdis* and *Bacteroides*, indeed encode *Bacteroides* Gene Ontology Pathways (**Supplementary Figure 5**), and not *Pdis*, further supporting that commensalism and interactions between these genera could be expected to be more frequent, compared to other genera, in CavFT lesions or other micro niches. This is supported by the fact that *Bacteroides* have been commonly isolated from CavFT in our studies^15,25^.

#### *The Pdis/B-fragilis* insert is present in mucosal rinsates from patient colonoscopies

The *B. fragilis* strain BFG539 corresponds to an isolate identified in human feces^29^. To assess the epidemiology in humans and the pathogenicity, tissue persistence, and transmissibility of CavFT-6^4^ in animal experiments, we use the location of such *B. fragilis* BFG539 fragment to develop a multiplex-PCR assay and expand the epidemiological utility of PCR and RT-qPCR protocol that we develop to amplify a *Pdis-*specific *rfbA* sequence^26^. Herein, we next developed primers to amplify two more bands, targeted at both 5’ and 3’ *Pdis/B-fragilis-*57,657bp (*CavFT/Bac*) junctions or insertion ends (**Figure 2E**). Once developed, to increase the external validity of our findings, we next conducted molecular epidemiological screening of bacterial isolates in other patients and various metagenome databases.

To better assess the prevalence of such *Pdis* CavFT-*B. fragilis* marker associated with gut mucosal persistence, we proposed to study the gut-associated presence of *Pdis* in the gut mucosa of ‘fasting’ humans, and not feces, because the presence in feces is confounded greatly with diet since gut microbiome abundance in feces may not reflect abundance in feces. To do so, we proposed to use colonoscopy as a strategy to study colonic mucosal rinsates from patients that were scheduled for colonoscopy at our University Hospitals Cleveland Medical Center under approved IRB protocols. In total, we obtained colonic rinsates from 80 patients: 50 patients with CD, 15 with ulcerative colitis (UC), and 15 from non-CD/healthy controls (HC). In short, patients that underwent screening colonoscopies had their digestive tract emptied prior to endoscopy with a 12-18h fasting and laxative ingestion protocol, which was deemed a better approach to study *Pdis* prevalence in mucosa associated surface niches, despite the influence of the laxative/fasting protocol.

Of note and as counterintuitively expected, the prevalence of *Pdis* was significantly lower in CD (and UC) rinsates compared to controls, as previously reported in studies with feces^30^, further supporting that *Pdis* is abundant in healthy individuals. Supporting the concept of regional clonality, multiplex PCR showed that 34% of all *Pdis*-positive patients also tested positive for the *Pdis/B-fragilis* marker sequence. However, against the hypothesis that CavFT would be more prevalent in CD, the proportion of CavFT-positive samples was higher in healthy controls, indicating that CavFT isolates are widely present in our community, but may not be pathogenic if humans do not have a predisposition to developing CD (**Figure 3A**). Although fundamentally counterintuitive to the proposition that CavFT *Pdis* is pathogenic, this epidemiological observation supports that this species could be probiotic in healthy individuals, but also illustrates that the identification of commensal bacteria deep inside CavFT lesions in surgical CD patients could represent the first proof-of-principle example of pathobiont biology where a commensal may drive inflammation or infection in chronic debilitating inflammatory digestive diseases. Presumably, this and other gut commensals, invade the gut wall in susceptible individuals, if such debilitating diseases allow for the formation of mucosal defects and CavFT lesions that enable the physical invasion of such commensals in a perpetual fashion where otherwise beneficial commensals cannot be cleared up from inside physical gut wall defects or microlesions.

**Figure 3.**
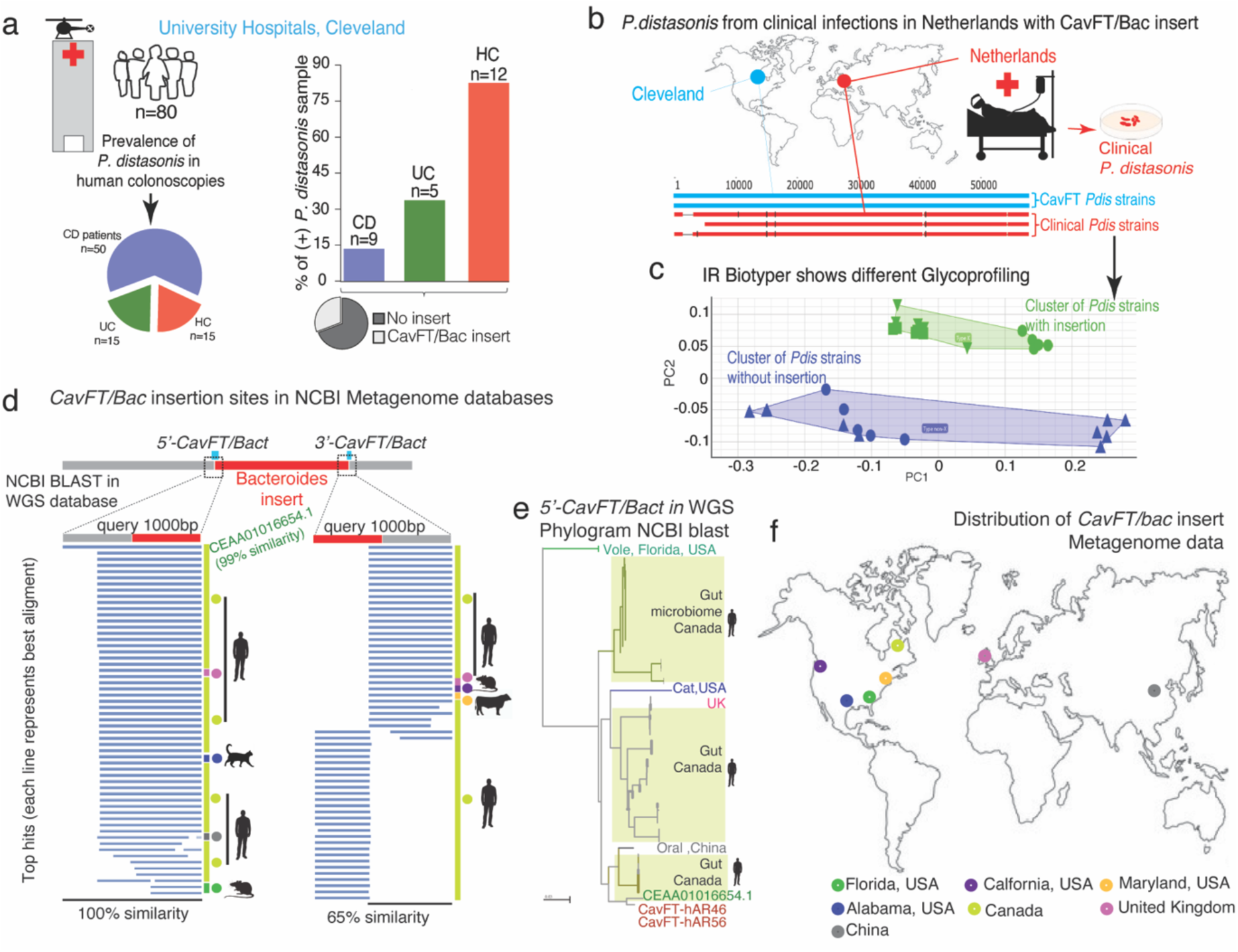
Molecular epidemiology of *CavFT/bac* insert in human and environmental metagenome. **a)** To epidemiologically assess the occurrence of the CavFT/*Bacteroides* insertion in other isolates, we conducted a multiplex-PCR primer test on 80 colonoscopy rinsates. A graphical representation to display the presence of the *rfb*A gene, which serves as an indicator for the presence of *P. distasonis*. Within the rfbA-positive samples, we employed a grayscale pie chart to illustrate the occurrence of the CavFT/*Bacteroides* cluster. **b)** Multiple Sequence Alignment reveals the presence of CavFT/Bac-insert in three clinically isolated *Pdis* medical strains from the Netherlands, along with two CavFT isolates. **c**) Surface glycoprofiling of isolates in panel 3b using IR Biotyper to compare the differences between the three Netherland isolates and ATCC 8503. Notice clusters in Principal Component Analysis (PCA). **d)** Multiple Sequence Alignment and similarity of published sequences compared to two 1000 bp regions from both 5’-CavFT/bac and 3’-CavFT/bac junction on Whole Genome Sequencing (WGS) data available on NCBI. Top NCBI BLAST hits of insert. **e)** NCBI-Blast Phylogenetic tree for 1000 bp regions from both 5’-CavFT/bac junction (**Supplementary Figure 7**). **f)** Global distribution of CavFT/Bac-insert best BLAST hits in WGS-metagenome database.

#### *The Pdis/B-fragilis* insert occurs in *P. distasonis* medical reference strains in the Netherlands

To further determine if such CavFT/B-fragilis insertion isolates were present in *Pdis* of clinical relevance to patients affected with serious medical infections, we collaborated with the anaerobic reference laboratory at the University Medical Center in Groningen, Netherlands^27^, where six representative *Pdis* isolates from patients with peritonitis/abscesses were shotgun sequenced to explore the genomic features of such insertion in Europe. Remarkably, three of the six isolates had virtually the same CavFT/*Bacteroides* insertion detected in our patients. Moreover, a whole cell extract glycolysis profile revealed that CavFT-positive isolates have a distinctive carbohydrate profile (**Figure 3B-C**, see PCA cluster vs ATCC-8503), supporting the concept that at least two clades of P distasonis exist for this species which has not been functionally illustrated before. The fact that *Pdis* can be divided into two groups or clades based on differences on carbohydrate surface composition reflect functional differences in both clusters that may imply differences in commensal or pathobiont opportunities for the clades.

#### *The Pdis/B-fragilis* insert internationally seen in digestive, not environmental, metagenomes

To determine if there is evidence of CavFT *Pdis* presence outside of the digestive tract (most genomes study gut-associated laboratory grown isolates), we next conducted a BLAST search for the CavFT/bac insert across the largest biorepository of metagenome databases (NCBI), by specifically selecting the Whole Genome Sequencing (WGS) database in NCBI, which contains sequencing annotated data from a diverse set of metagenome studies conducted in numerous locations. Considering that metagenomics commonly uses shotgun metagenomics to generate large libraries of small-sized fragments for each sample, we opted to select smaller sequence regions to explore the hypothesis that such emerging CavFT *Pdis* strains are not distributed worldwide. Thus, we selected two 1000 bp sequences to encompass the junction regions for both the 5’-CavFT/bac and 3’-CavFT/bac insert sites (500bp on the *Pdis* side, and 500bp on the *B-fragilis* side) as shown in **Figure 2D** for this BLAST WGS analysis. Our analysis surprisingly revealed almost 100% similarity in the 5’-CavFT/bac region for only one metagenome sample in NCBI corresponding to the human gut metagenome of an individual involved in an antibiotic screening study launched in 2016 in the province of Quebec, Canada^31^ (**Figure 3D-E, Supplementary Figure 7**). Of interest, the remaining top BLAST hits occurred across other participants in the same study with only approximately 100-40% similarity (±14% SD; 20% less similar on the *Parabacteroides* site than on *Bacteroides* site). Or interest, the 3’-CavFT/bac region showed a 70% similarity in Canada. Intriguingly, during our investigation, we also identified the presence of the CavFT/Bac insert in China from a sample of human oral metagenome (**Figure 3F**). Remarkably, we were able to identify better matches for the 5’-CavFT/bac insert site, than for the 3’-CavFT/bac site. Despite these findings, we were not able to detect this CavFT/bac insert in environments other than the digestive tract, through this analysis when we filtered our analysis to include various aquatic, terrestrial and unanimated environments. To verify such observation, we then used the Joint Genomics Institute metagenome databases and (JGI-IMG/MG) tools to search for an additional collection of primarily environmental metagenomes and host-metagenomes but could not identify find any significant hits.

#### *CavFT P. distasonis* is innocuous in healthy mice but worsens CD-like ileitis in SAMP mice

To assess the pathogenic potential of CavFT-46^4^ in experimental CD, we next conducted colonization studies using the well-established mouse line, SAMP1/YitFc, which was discovered in 1980^16^, and remains the only mouse model known to develop a form of CD ileitis that structurally resembles characteristic lesions observed in humans with ileal CD^2, 16, 32, 33^. While SAMP mice develop CD-like ileitis under germ-free (GF) conditions, indicating that chronic CD-like inflammation in these animals does not strictly require bacteria^30^, we found that colonizing 20-week-old GF-SAMP mice with CavFT-46 for 6 weeks worsened the severity of cobblestone ileal lesions (**Figure 4A**). Since we hypothesized that the susceptibility of host mice to developing ileal inflammation facilitates colonization by CavFT-46, we conducted 2-week colonization experiments with GF-SAMP and control GF-C57BL/6J (B6, ileitis-free/healthy) mice to determine if CD-predisposed mice favored the colonization of CavFT-46 compared to healthy control mice. In support of this possibility, CavFT-46 was detectable and more abundant on ileum tissues from SAMP mice compared to healthy mice, while no substantial differences were observed in fecal samples (**Figure 4B**, see RT-qPCR data). By using fluorescent in situ hybridization (FISH) and *Pdis* specific primers, we verified the presence of CavFT-46 within the ileum of SAMP mice confirming that bacteria penetrate the gut wall of CD-like ileitis-affected mice (**Figure 4C**).

**Figure 4.**
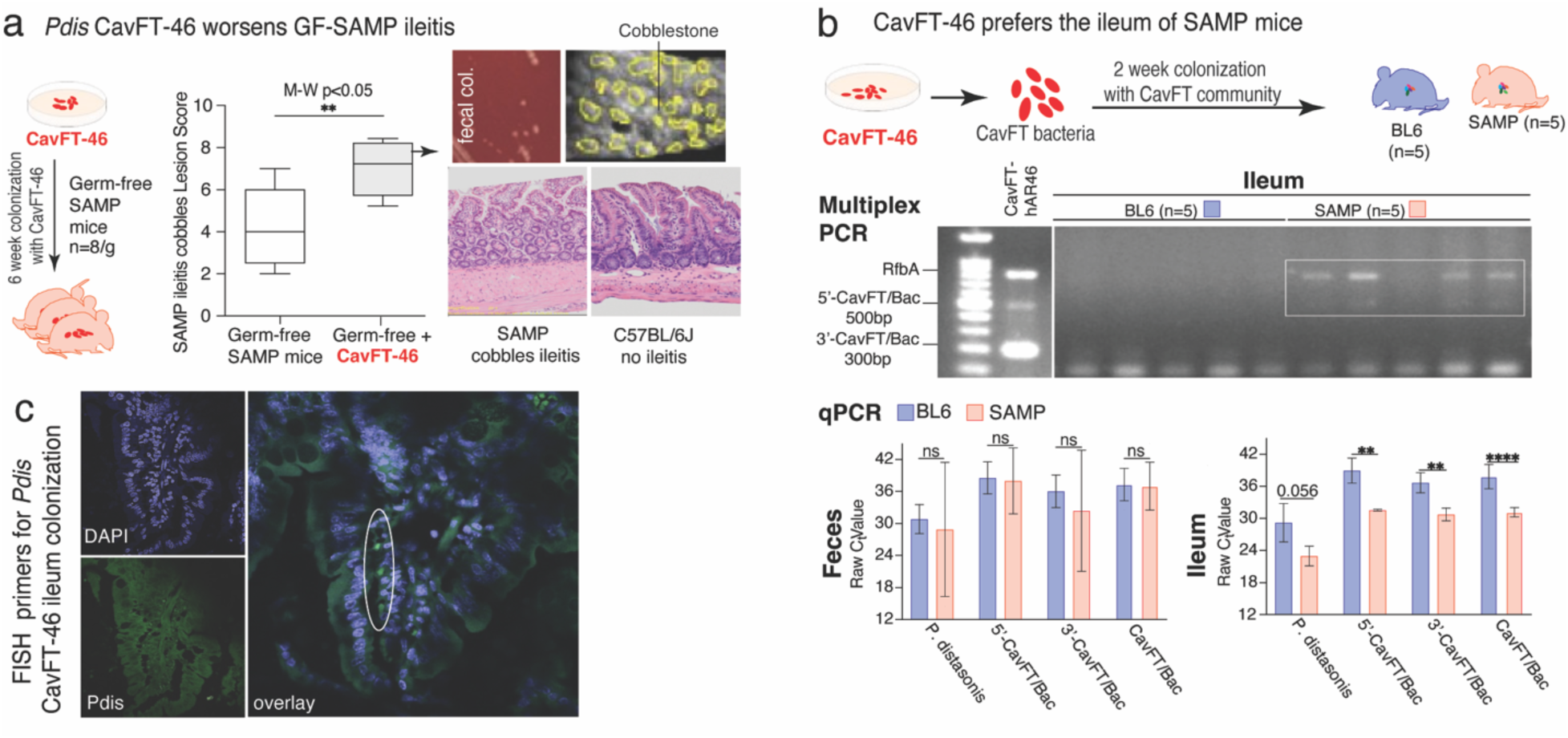
Experimental invasion of CavFT bacteria in the ileum of CD-prone SAMP mice. **a)** Overview of 6-week colonization experiment of CaVFT-hAR46 for six weeks in germ-free (GF) SAMP mice as model of CD ileitis. Inset pictures, TSA agar showing pure *Pdis* colonies from feces of GF-SAMP mice at the end of study; ‘*en face*’ view of SAMP ileum mucosa with juxtaposed cobblestone lesions (dashed circles) affecting 80% of ileal mucosa; H&E histology of ileum in SAMP and C57BL/6J mice showing typical abnormal villi and serosa thickening in SAMP mice. **b)** Overview of colonization experiment of CavFT-hAR46 in germ-free (GF) SAMP mice to assess potential for bacterial invasion and adaptation in the ileum tissue. Inset, gel electrophoresis and qPCR on the ileal tissue of colonized SAMP and C57BL/6 control mice. Notice that CavFT-hAR46 bacteria invade and are detectable in the SAMP ileum, not C57BL/6 controls**. c)** Ileal sections from SAMP mice supplemented with *Enterococcus faecium, Klebsiella variicola*, and CavFT-hAR46 were subjected to hybridization using probes specific to *P. distasonis* (visualized in green). The cell nuclei were visualized using DAPI staining (appearing in blue).

#### Immortalized myeloid cells from SAMP mice reveal host susceptibility to succinate and *E. coli*

Previous studies suggest *Pdis* has potential as both an anti-cancer agent and probiotic^27^, but conflicting reports have been published with regard to pathogenic potential. The mechanisms underlying *Pdis*’ switch from commensal to pathobiont, especially in Crohn’s disease (CD), remain unclear. Recent findings highlight differences in the O-antigen operon, vital for LPS production, between *Pdis* and *E. coli*. In vitro, *Pdis* exhibited impaired LPS production and produced a lower inflammatory response in murine macrophages when compared to *E. coli*, further underlining its paradoxical non-pathogenic behavior^15^. We hypothesized that a specific metabolite-bacteria interaction could drive *Pdis* pathogenicity in CD^34–36^, especially when the host is susceptible to CD and has abnormal bacterial clearance, as in SAMP mice^37^. To test this hypothesis, we used target metabolomics of the glycolysis pathway in CavFT-hAR46 using C13-glucose and found elevated succinate production compared to other Krebs’s cycle metabolites (**Figure 5A**), as well as to the metabolome of human colon cells (Caco2).

**Figure 5.**
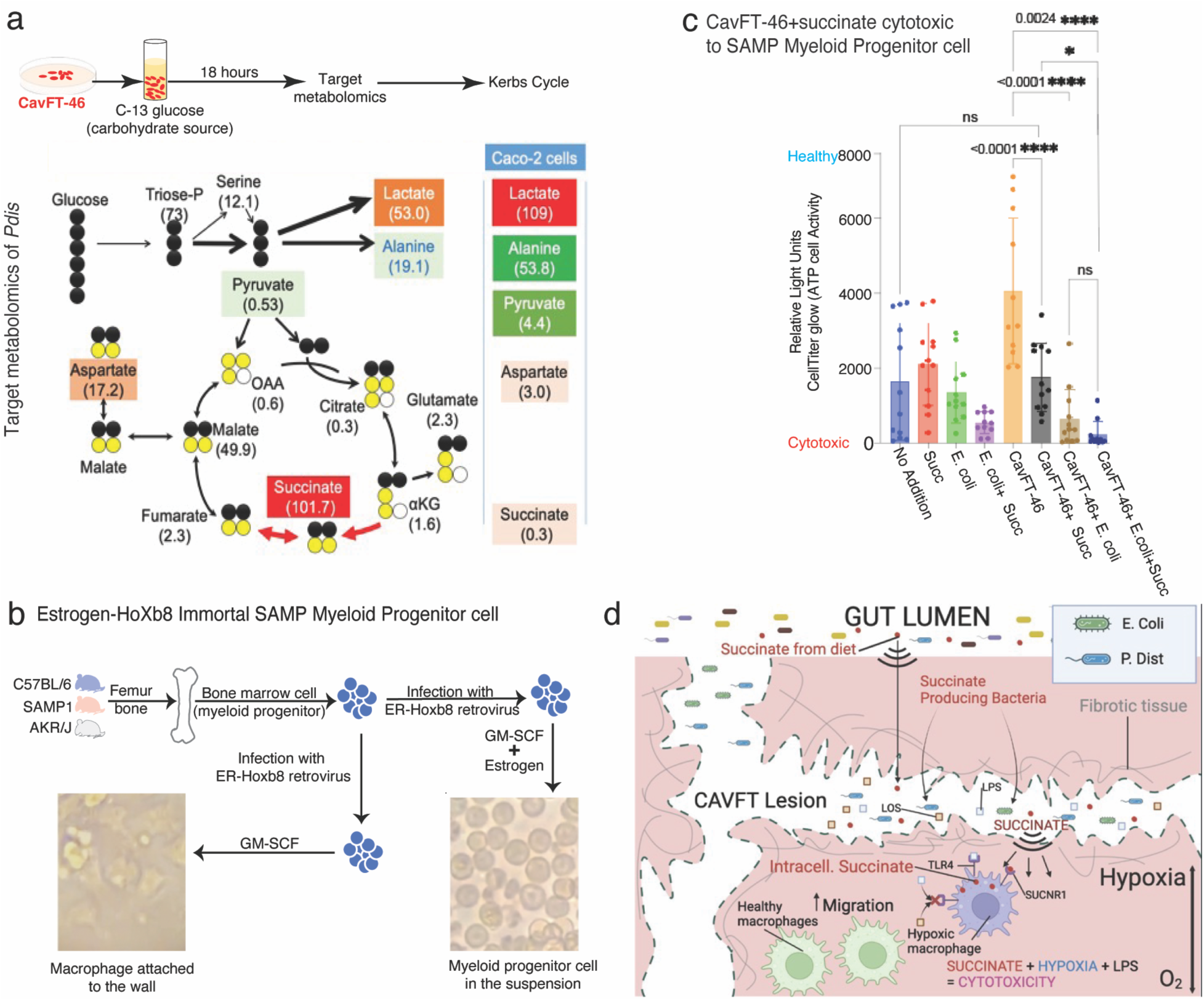
Model of succinate mediated pathogenicity of *P. distasonis* in immortalized myeloid cells from mice prone to CD-like ileitis. **a)** Target metabolomics with C13-glucose shows *P. distasonis* results in rich production of succinate anaerobically. A virtually identical succinate-rich profile has been identified in other isolates from CavFT that belong to *Bacteroides* and *Alistipes* spp.^25^ (data not shown). **b)** Overview of experiments to generate our estrogen-HOXB immortal myeloid progenitor cells from SAM/YitFc and control mice. c) *P. distasonis* is highly toxic in the presence of succinate and *E. coli* to SAMP/YitFc immortalized bone marrow-derived cells. ANOVA, Bonferroni pairwise statistics. We measured cell toxicity using Cell titer glow (Promega) n=12 replicates/conditions. **d**) Hypothetical interactions of succinate-producing commensal bacteria (*P. distasonis*) and *Enterobacteriaceae* (*E. coli*) co-habiting in CavFT and its dependence on tissue hypoxia which promotes host succinate and inflammation.

Since succinate can trigger immune-mediated inflammatory responses^38^, we next developed a novel panel of IBD-related immortalized cell lines derived from SAMP mice (prone to IBD) along with C57BL/6, AKR/J (control mice) (**Figure 5B**). Utilizing *in vitro* methods with real-time and time-lapse cell imaging (Incucyte system), we discovered that SAMP cells exhibit heightened sensitivity to succinate exposure, unlike the C57BL/6 and AKR/J myeloid progenitor cell lines. This was evident through a red cell count, and quantifying the presence of dead cells using indirect cell-pump functionality and cell viability using propidium iodide. This remarkable finding for the first time illustrates that SAMP-ileitis susceptibility scales down from the 100% penetrant animal phenotype to an immune cellular functional phenotype of susceptibility to a glycolysis metabolite (succinate) that is commonly found in numerous gut commensals (*Bacteroidota*) and that is also produced in abundance by host cells in the mitochondria under hypoxic conditions. This pattern of cellular susceptibility could explain why *Pdis* CavFT could be pathogenic to CD individuals, and not to healthy humans, which presumably would not have such a cellular susceptibility to succinate.

With this remarkable CD-like ileitis cellular phenotype from SAMP mice, we next tested interactions that could include bacteria present in CavFT. We thus quantified the effect of several concentrations of succinate, with and without the presence of CavFT *Pdis* and other bacteria on a clone of such immortalized SAMP myeloid cells *in vitro*. For this assay, we switched to assessing mitochondrial health and ATP abundance in viability assays by CellTiter-Glo using 356-well formats

Remarkably, SAMP myeloid cells displayed a very potent reduction in ATP production when succinate *Pdis* and *E. coli* were combined, while of great interest heat-treated extracts of *Pdis* CavFT-46 alone were highly beneficial for cell protection and ATP enhanced production. This finding could potentially explaining how a set of combinatorial factors facilitate the transition of an otherwise beneficial commensal organism from being ATP promoter to becoming a more toxic and pro-inflammatory pathobiont in a CD-prone host if co-habiting with other bacteria like *E. coli*, a hypothesis we earlier proposed when examining the operon integrity and DNA sequence evolution for this and other *Bacteroidota* species^15^.

Additional mechanistic studies are underway to further explore this hypothesis (**Figure 5C**). Mechanistic roles of succinate in the pathogenicity of CavFT bacteria remain unclear, but in the case of gram-negative bacteria, LPS amplifies succinate (SUCC) uptake by macrophages, the expression of SUCNR1, and the production of succinate within the cell. Elevated intracellular SUCC levels lead to an increase in inflammatory proteins, initiating cascades that result in heightened interleukin-1β (IL-1β) levels^18, 38–42^. It is known that the expression of IL-1β protein induced by LPS increased in hypoxia condition^43^. The presence of CavFT likely initiates hypoxic cascades as cells approach it, amplifying an inflammatory cascade at the site where bacteria thrive. This may perpetuate a feed-forward cycle of chronic inflammation which herein we proposed a succinate-susceptibility genetic mechanism as a possibility to partly explain the CD-ileitis dominance in SAMP mice compared to the resistance of the parental AKR/J mouse line to any form of inflammatory bowel disease susceptibility or microbiome-driven transmissibility of the SAMP-IBD phenotype as we showed earlier in long-term SAMP1/YitFc:AKR/J cohabitation studies^44^(**Figure 5D**).

#### Transmission of *P. distasonis* CavFT46 among SAMP mice is absent in MPO-deficient BL6 mice

To strengthen our hypothesis that the host predisposition to CD-like phenotype facilitates the colonization of SAMP mice by CavFT *Pdis*, we conducted *Pdis* transmission experiments with groups of conventional SPF-SAMP mice harboring regular SPF microbiota, and with germ-free mice a second mouse model where GF-SAMP mice were cohoused with mice pre-colonized with CavFT-46. Of importance, experiments proved that not every single exposure to CavFT-46 leads to successful colonization by *Pdis*. Upon 7-days of cohabitation of mice with a littermate colonized and gavaged with CavFT-46 *Pdis,* transmission of *Pdis* from the colonized/gavaged mice was observed in some SAMP mice when they were raised under SPF-microbiome conditions (n=1/3, 33.3%), while transmission was rapidly observed in all exposed SAMP mice when reared under germ-free conditions (n=3/3; 100%; **Supplementary Figure 8**) supporting that *Pdis* transmissibility may occur via cohabitation and the fecal-oral route (presumably grooming or coprophagia), and that the resident gut microbiome in SPF mice is protective against *Pdis* colonization in some cases.

Since there are no other models of CD-like ileitis that resemble cobblestones lesions in humans^16^, we next determined if a more genetic form of susceptibility to bacterial superinfections due to defects in innate immunity could increase the susceptibility of mice to CavFT-46 *Pdis* colonization and fecal-oral natural transmissibility. Thus, to determine if a deficient immune genotype, facilitated *Pdis* colonization, we used homozygous SPF myeloperoxidase deficient C57BL/6J (B6^MPO-/-^) mice^45^. In short, B6^MPO-/-^ mice were studied in triplicates of 4 mice/cage, with one of the cohoused mice being gavaged with CavFT-46 *Pdis* (10^7-8^ CFU) every 24-48h. Of interest, after 7 days of repeating gavaging and cohabitation, none of the mice gavaged, or cohoused with the gavaged mice, were positive to CavFT-46 *Pdis* either by our multiplex rfbA-PCR testing or culture. These results indicated that the gut microbiome and possible the genomics of the C57BL/6J host are protective against colonization, in a manner that is independent of MPO-deficiency, the ensued susceptibility to bacterial superinfection, and presumably the gut microbiome since all mice were tested at the same time and bred/raised in our SPF animal facility (4/6 in SAMP, vs 0/12 B6^MPO-/-^, Fisher exact P=0.0049). This is an interesting finding that requires further investigation.

For environmental survival, we found that CavFT-hAR46 did not die when left to dry on blood agar at room air and 23°C for 9 days. Supporting that *Pdis* is amenable to environmental transmission, we determined that pure cultures of *Pdis* CavFT-46, despite being a strict anaerobe for growth, can survive aerobic incubation (room air) and dehydration (left on tryptic soy agar surface) and be recoverable by replanting the colonies on fresh pre-reduced TSA agar. To ensure proper anaerobic conditions, the TSA agar was incubated in the anaerobic chamber for 3 hours (pre-reduced to allow for oxygen effusion) before plating the colonies. Of notoriety, recovery was not possible if subculture occurred on non-pre-reduced agar (strict anaerobiosis for regrowth)^2^. During active growth in liquid medium (BHI), exposure to room air for 20 minutes transiently inhibited CavFT-hAR46 grow.

## Discussion

Genetically, this study represents the first complete genome comparative analysis of *P. distasonis* derived from gut wall microfistulous lesions from CD patients, and a mechanistic commensal model of CD complications where pathogenicity is host-dependent on the synergistic presence of the commensal, succinate and *E. coli*. This model could explain sustained inflammation and poor response to antibiotics since these principles of susceptibility are common among other commensals (*Bacteroidota*) that abound in such micro-niche communities.

The role of inflammation in CD is prominent in medicine that consensus guidelines^46, 47^ instruct the use of immunosuppressants, discouraging the use of antibiotics. This study highlights the potential role of gut commensals on CD inflammatory micropathologies. We do not believe that CD is caused by bacteria, since CD-like ileitis occurs in SAMP1/YitFc mice raised under germ-free conditions^2^, supporting that host-susceptibility is the main driver of CD (not bacteria). Fecal matter microbiota transplants from CD patients in remission however worsens the ileitis in GF-SAMP mice^3^, but not all CD patients have pro-inflammatory microbiotas.

Results from the present study are highly significant because it could help explain how presumed beneficial commensals may turn non-intentionally into pathogenic microorganisms in the gut wall of susceptible hosts. We found that succinate susceptibility could promote failure of immune cell functions in a matter that is theoretically proportional to the amount of succinate produced, the cohabitation with other commensals or gram-negative enterobacteria and possibly distance from site where bacteria thrive^2^. Despite a major advancement in our understanding of this proof-of-concept, there are numerous open questions with regards to the genetic mechanisms that in the case of SAMP ileitis and humans that could enable the presence and long-term survival of *Pdis* in the gut wall. Also uncertain are the precise factors from the CavFT *E. coli* used in this study (CavFT-359 *E. coli*, such as LPS^15^).

Infection and transmission experiments with the SAMP1/YitFc mice (prone to CD-like ileitis), demonstrate that these CavFT-derived *Pdis* isolates are ecologically adapted to invade the ileum wall and worsen the ileitis of mice presumably due to their susceptibility to CD, but not in healthy control mice. Since *Pdis* has many anti-inflammatory properties and lack LPS formation due to O-antigen operon fragmentation^15^, the generation of immortalized mouse myeloid cell lines represent a novel assays that ecologically and phenotypically could be extrapolated to humans to assess the role of such CavFT isolates directly in human immune myeloid cells. Of interest, in this assay system, *Pdis* extracts alone promoted cell ATP activity^48^, on the same cells that were highly susceptible to succinate/*E coli* co-toxicity. How the same metabolite can be so beneficial alone but be toxic in other scenarios remains unknown.

In conclusion, our genomic, metabolomic and colonization studies on *Pdis* reveal for the first time that the CavFT bacteria isolated from these CD patients may be both uniquely adapted to thrive in ileal microlesions and potentially transmissible across susceptible hosts in the community. Our mouse study indicate that *Pdis* may not be pathogenic in healthy individuals. However, *Pdis* isolated from CavFT from CD-patients worsen the ileitis in GF-SAMP mice indicating a contributing role in CD microlesions. This establishes a foundation for further exploration, linking *Pdis* to CD disease complications fulfilling the Koch’s postulates^49^. The reason for the SAMP/human-dependent susceptibility remains unclear. Given the genomic clonality and ecological uniqueness, it is possible these CavFT *Pdis* isolates represent a gut-wall adapted subspecies based on genetics/ecological criteria^4^. Herein, we propose the designation of *P. distasonis* nov. subspecies ‘*cavitamuralis’* to reflect the ecological ability of some commensals to be genetically distinct and able to thrive in the gut wall or associated microlesions (LPSN-distinctive; *‘cav’ for cavita*/cavity/CavFT; and “*muralis*” for “*intramuralis”* to reflect gut-wall adaptation^50^). The host-driven mechanisms that explain the immune susceptibility to such a combination of factors are expected to be very complex but the role of succinate in human IBD^18, 38, 42^ and SAMP ileitis is currently investigated in CD.

Our data supports the proposition of a host-conditional succinate cytotoxicity commensal model of CD complications where succinate-producing bacteria (in *Bacteroidota,* not only *Pdis*) in CD microlesions may perpetuate inflammation and pathology by impairing immune cells in succinate-susceptible hosts. Our study and others further indicates that *Pdis* species could be potentially beneficial (probiotic) in healthy humans^27^; however, our findings in CD raise concerns about the use of this or other uncharacterized commensal succinate-producing bacteria in CD patients.

### Online Methods

#### Sample collection and Culturing

This strain was obtained from CavFT, which was found in the bowel and surgically removed from a severe Crohn’s disease patient^4^. The bowel sample was dissected and cultured under anaerobic conditions using the previously described method^4^. Through 16S RNA Sanger sequencing and MALDI-TOF (matrix-assisted laser desorption ionization–time of flight), the cultured strain was identified as *Parabacteroides distasonis* (Taxonomic ID 823). DNA was isolated from the cultured bacteria using the Qigen MagAttract HMW DNA kit (Qiagen). The clinical sample was obtained from the NIH Silvio O. Conte Cleveland Digestive Disease Research Core Center in Cleveland, Ohio, USA, in accordance with proper consent and approval obtained from the patients’ protocols, which were approved by the institutional review boards of University Hospitals Cleveland Medical Center and Case Western Reserve University.

#### Genome sequencing and processing

The genomes were sequenced utilizing Pacific Biosciences (PacBio) reagents obtained from DNA Link, Inc. (South Korea), and the library was generated using the SMRTbell template prep kit 1.0 (product number 100-259-100; PacBio). The validation of the library was performed using an Agilent 2100 bioanalyzer, and SMRT (single-molecule real-time) sequencing was carried out on the PacBio Sequel platform using P6-C4 chemistry (DNA sequencing reagent 4.0). All software was used with default parameters unless specified. The Flye 2.56 software was utilized for de novo assembly. For CavFT-hAR56, a total of 4,951,235 bp was distributed across two contigs, which were error-corrected using reads (contig N50, 4,946,368 bp, for a total of the 2-overlapping contig, a total length of 4,951,235 bp) with an average sequencing coverage of 183X. For CavFT-hAR46 re-sequence control, a total of 4,952,355 bp was distributed across two overlapping contigs, which were error-corrected using reads (contig N50, 4,944,140 bp, 2 contigs with a combined length of 4,952,355 bp) with an average sequencing coverage of 168X.

#### Genome analysis of CavFT-hAR56 with other *P. distasonis* Genome

For contextualization, we utilized the genome database available in Bacterial and Viral Bioinformatics Resource Centers (BV-BRC) for computerized annotation and general comparison of CavFT sequenced strains with other available strains of *Parabacteroides distasonis*.

#### Construction of phylogenetic tree of different strains of *P. distasonis*

We also utilized the Bacterial and Viral Bioinformatics Resource Centers (BV-BRC) for phylogenetic tree construction. Specifically, we employed the PATRIC Codon Trees pipeline, which utilizes 100 randomly selected protein and DNA sequences from PGFams (PATRIC’s global Protein Families) for alignment and tree construction using the RaxML program. Both amino acid and nucleotide sequences were used in the phylogenetic tree construction. The tree branches were visualized using iTOL software.

#### Genome alignment to show genome rearrangement and a blast of ATCC 8503, CavFT-hAR46 and CavFT-hAR56 with each other

We used the Mauve software for the genome alignment of CavFT-hAR46, CavFT-hAR56, and the reference genome ATCC 8503.

#### PCA plot of Genome alignment of ATCC8503 and 82G9 strain with 12 different strains of *P. distasonis*

The whole genomes of ATCC 8503 and 82G9 were aligned with 12 different strains of *P. distasonis* using BV-BRC between the length of the nucleotide align together and the number of times they align together. Mauve software was used for genome alignment, which provided information about nucleotide sequence positions aligned together in the two genomes. Statistical differences among the number of nucleotide sequences in the genomes of ATCC 8503 and 82G9 aligned together with the 12 different strains were identified through Principal Components Analysis (PCA) biplot. PCA plot formation was constructed using GraphPad Prism 9 software.

#### Genome Alignment analysis of CavFT-hAR(46 & 56) with different strains of *Bacteroidota* to find the transfer of genetic material between two *Bacteroidota*

We utilized BV-BRC, which employed Mauve software for genome alignment, providing information about nucleotide sequence positions aligned in both genomes. The total nucleotide sequences aligned together were summed to calculate the percentage of genetic material transferred between them.

#### Construction of 16s and 23s RNA Phylogenetic tree

The phylogenetic tree of 23S and 16S RNA was constructed using maximum likelihood. We utilized the IQ-TREE^51^ webserver with default parameters and 1000 bootstrap replicates to generate the tree. The phylogenetic tree branches were visualized using the iTOL software^52^.

#### BlastX and metagenomics analysis of 57657bp CavFT/*Bacteroides* insert of CavFT-hAR56

We downloaded the 57657bp sequences of CavFT/*Bacteroides* insert from NCBI, and then split it sequences into 250bp long sequences fragments using online software (https://www.bioinformatics.org/sms2/split_fasta.html). We performed BlastX and metagenomics analysis using these 250bp long sequences fragments.

#### Methodology IR Biotyper

*Parabacteroides distasonis* isolates were subcultured on Brucella Blood Agar (Mediaproducts, Groningen, the Netherlands), at 37 °C in an anaerobic atmosphere for 48 hours. Bacterial cells were harvested by suspending a 1 µl loop full of bacterial biomass from the confluent part in a solution of 50 µl 70% ethanol and 50 µl of water in Eppendorf with steel cilinders (IR Biotyper® kit, Bruker Daltonics). After thorough mixing, 15 µl of the suspension was pipetted on a spot of the silicone target (Bruker Daltonics, Bremen, Germany) in 5 technical replicates. The target was left to dry in an incubator at 37 °C, 5% CO_2_ for 10-20 minutes. Spectra were acquired and processed using the IR Biotyper ® system (Bruker Daltonics GmbH & Co. KG) and the OPUS v8.2 software (Bruker Optics GmbH & Co. KG, Germany) in transmission mode in the spectral range of 4,000-500 cm-1 (mid-IR). Spectra processing and visualization were performed with the IR Biotyper ® Client software V4.0 (Bruker Daltonics GmbH & Co. KG) using the default settings. Spectra were smoothened using the Savitzky-Golay algorithm over nine data points, whereafter the second derivative of the spectra were calculated. The spectra underwent splicing within the 1,300-800 cm-1 (carbohydrates absorption) range and were subsequently vector-normalized, aiming to address the preparation-induced variability in biomass and absorption. For each run, the Infrared Test Standard 1 and 2 (provided in the IR Biotyper® kit) were applied. All spectra were acquired by undercalculating a background spectrum between each sample/control measurement.

#### IR Biotyper data analysis

Multivariate exploratory data analysis was performed by the IR Biotyper Client v4.0 software. Principal Component Analysis (PCA) Linear Discriminant Analysis (LDA) with Isolate ID as target group were used to visualize the data.

#### CavFT/Bac insert search in Metagenome database (WGS)

To identify the Bac/CavFT insert gene in microbiome, we conducted NCBI Blastn searches using whole-genome shotgun contigs (wgs) for the 1000 bp regions from both *5’-CavFT/bac* (14683-15683 for CavFT56 & 2198759-2199759 for CavFT-46) and *3’-CavFT/bac* (72440-73440bp for CavFT-56 &. The organisms included in the analysis were sourced from the human microbiome (Taxid 646099), gut microbiome (Taxid 749906) and human oral microbiome (Taxid 410658).

#### 3D-stereomicroscopic analysis of GF-SAMP/YitFc mice

We conducted an experiment using two groups, each group consisting of 20-week-old eight GF SAMP1/YitFc mice (prone to IBD). One group underwent a 6-week colonization with CavFT-hAR46 bacteria, while the other group remained germ-free without any bacteria. After euthanizing the mice, we harvested ileum tissues and assessed lesions using 3D-stereomicroscopic analysis, as previously described^2^. In summary, we started by focusing on the distal ileum, focused on a 1-cm ileum field, and initiated the scoring of cobblestone-like structures.

#### Multiplex PCR for validation of CavFTs strains and 80 colonoscopy rinsates

A mixture called the master mix was prepared, which consisted of ThermoFischer Scientific DreamTaq Green PCR Master Mix (1x), RNase-free water, and sets of primers for the ‘CavFT/*Bacteroides* exchange cluster’.i.e 5’-CavFT/bact(F-5ʹ-ATATGGAAACTAAATTTACATGGATACC-3ʹ;R-5ʹ-GAACATATTTTTCCGGATTAACGTAGA-3ʹ),3’-CavFT/bact (F-5ʹ-TTGTTCCTTATTTGTTTCTCTATTAAAACT-3ʹ; R-5ʹ-GGGGTCGAGTTGGGTACCGAGGATATC-3ʹ), and *P. distasonis* validation *rfbA* Primer (F-5ʹ-CCGCTTGTATCCGATCACT-3ʹ; R-5ʹ-AAATACTGGCCGTACTGATTCTT-3ʹ)^26^ to validate *P. distasonis* species and 3 µL of *P. distasonis* strains DNA. The DNA was then amplified using the hot start 95 °C, 5 min and 45 amplification cycles; 95 °C, 30 s; 55 °C, 30 s; 72 °C, 45 s, followed by 4.0°C indefinitely using the Applied Biosystems Veriti Thermal Cycler. The resulting amplified products, along with an exACTGene 100 bp DNA ladder (10 µL), were then subjected to agarose gel electrophoresis (1%) containing ethidium bromide (2 µL) at 100 V for 30 minutes. The gel was visualized under UV light.

#### qPCR showing CavFT-hAR46 dynamics in Humanized IBD and non-IBD mouse models

We conducted an experiment on (3 mice/group) of age-matched and sex-matched 7-week-old GF SAMP1/YitFc mice (Prone to IBD) and C57BL/6 (Control). All mice were individually caged using our GF-grade nested isolation (NesTiso) caging system (Cyclical Bias’ in microbiome research revealed by a portable germ-free housing system using nested isolation.) and maintained on nonedible Aspen bedding and the mice were orally administered with the community of bacteria (*P. distasonis* CavFT-hAR46, *E. coli*, *Klebsiella variicola*, *Alistipes finegoldii*, *Enterococcus* sp, *Pediococcus sp.*) and the mice were sacrificed after 14 days to collect the tissue sample. The DNA was isolated from the ileum using the Qigen DNA extraction kit and qPCR was performed with five different primers Universal (F-5ʹ-TGGCTTAGTTTACCAGAGTCATA-3ʹ; R-5ʹ-GGACTACCAGGGTATCTAATCCTGTT-3ʹ), 5’-CavFT/bact (F-5ʹ-ATATGGAAACTAAATTTACATGGATACC-3ʹ; R-5ʹ-GAACATATTTTTCCGGATTAACGTAGA-3ʹ), 3’-CavFT/bact (F-5ʹ-TTGTTCCTTATTTGTTTCTCTATTAAAACT-3ʹ; R-5ʹ-GGGGTCGAGTTGGGTACCGAG GATATC-3ʹ) and *P. distasonis* (F-5ʹ-CTTGCCACTAGTTACTAACA-3ʹ; R-5ʹ-CCCTGTCGCCAGGTG-3ʹ) primers. 1 ul of DNA is used in a final reaction volume of 20 μl. qPCR amplification of genomic DNA samples was performed using clear 96-well plates and the Roche 480 LightCycler SYBR Green run template settings (hot start 95 °C, 5 min and 45 amplification cycles; 95 °C, 15 s; 53 °C, 30 s; 72 °C, 30 s).

#### Transmission of CavFT-hAR46 in SPF-SAMP and GF-SAMP/YitFc

We conducted an experiment on 20-week-old 4 SPF-SAMP1/YitFc mice (prone to IBD) and 3 GF-SAMP1/YitFc mice, which were age-matched and sex-matched. One mouse from each group was orally administered with CavFT-hAR46 and then co-housed with other mice. After seven days, feces were collected, and DNA was extracted using the Qiagen DNA extraction kit. Multiplex PCR was performed, as discussed earlier, to check the transmission of CavFT-hAR46 from one mouse to another.

#### FISH to localized and identification of CavFT-hAR46 in the Ileum

To localize and identify CavFT-hAR46 in the ileum of the mice gavaged with *Enterococcus faecium*, *Klebsiella variicola*, and *P. distasonis* CavFT-hAR46, we employed FISH analysis. Sections of Carnoy’s fixed paraffin-embedded colonic tissue (6 μm thick) were mounted on Probe-On Plus slides (Fisher Scientific, Pittsburgh, Pa.). These sections were then deparaffinized using xylene (three washes) and a series of graded alcohol solutions (100%, 95%, 70%). Subsequently, the sections were treated with lysozyme (15 U/μl in 1% PBS) at 37°C for 20 minutes, followed by two rinses with PBS and water. For FISH hybridization, the sections were incubated overnight at 56°C with specific probes for *P. distasonis* labelled with (Alexa-657 5’-CCGCCTGCCTCAAACATA). After hybridization, the slides were washed in hybridization buffer without SDS at 58°C for 20 minutes, followed by rinses in 1% PBS and distilled water. Subsequently, the slides were dried at 46°C for 10 minutes and mounted using the ProLong antifade Dapi kit (Molecular Probes Inc., Eugene, OR). The sections were examined using an Olympus FV1200 Confocal Microscope. Images were captured using the Olympus DP-7 camera or Zeiss Axiocam, respectively.

#### Targeted metabolomics of CavFT-hAR46

MLE12 bacterial cells were treated with 20 millimoles of [U-13C] glucose for 18 hours. After this exposure, the cells were collected, and the changes in metabolic activity were studied by observing how the 13C labeled carbon was integrated into different metabolites. Following the incubation period, a 50 µl culture media was taken and stored for analyzing the initial compounds used in making cellular components. The bacterial samples were preserved by freezing them at -80°Celsius until they were ready for further examination. The method described by van Dijk, TH, and others in JBC, 2001 was adjusted and used to determine the enrichment of 13C-glucose. The samples were mixed and left on ice for 30 minutes. Following this, the samples were spun in a centrifuge at 4°C for 10 minutes at 14,000 rpm, the ethanol was moved to a GC/MS vial and evaporated to dryness in a SpeedVac evaporator. The glucose present was transformed into its pentaacetate derivative by a chemical reaction involving 150 microliters of acetic anhydride in pyridine (in a 2:1 ratio of volume) at a temperature of 60°C for 30 minutes. After the sample was dried, the resulting derivative of glucose was dissolved again, this time in 80 µl of ethyl acetate, and then placed into the GC/MS instrument. Masses 200 and 205 were observed and recorded after samples were injected in duplicate. Glucose (precursor enrichment) was calculated as (205) / (200+205).

#### Metabolite extraction

Add 1 ml (80% methanol), and the cell extract was vortexed and sonicated for 10 minutes with an ultrasonicator for 30 seconds on, 30 seconds off, alternating. Following that, the cells were pelleted by centrifugation at 4°C for 10 minutes at 14,000 rpm. The supernatant was transferred to GC/MS vials and dried under a gentle stream of nitrogen. By adding 10 l of 1 N NaOH plus 15 µl NaB2H4 (prepared as 10 mg/ml in 50 mM NaOH), keto- and aldehyde groups were reduced. After mixing, samples were incubated at room temperature for 1 hour before being acidified with 1 N HCl (55 µl) by dropping slowly. Following that, samples were evaporated to dryness. Then, to precipitate the boric acid 50 µl of methanol. Internal standard was added (10 µl of gamma-hydroxybutyric acid, 0.1 mg/ml). After drying, samples were to reactions with 40 µl of pyridine and 60 µl of tert-butylbis(dimethylsilyl) trifluoroacetamide with 10% trimethylchlorosilane (Regisil) TBDMS at 60°C for 1 hour. TBDMS derivatives were produced and injected into GC/MS.

#### GC/MS conditions

Agilent 5973 mass spectrometer equipped with a 6890 gas chromatograph was used for analyses. All assays were conducted using an HP-5MS capillary column (60 m × 0.25 mm × 0.25 μm) with a helium flow rate of 1 ml/min. Electron impact ionization (EI) was used to analyze the samples in the Selected Ion Monitoring (SIM) mode. 10 msec was the ion dwell time that was set.

#### Calculations

To determine the Fractional metabolic flux a relationship between precursor ([U-13C]glucose) to product ([13C]-labeled metabolites) was applying. Fractional metabolic flux was calculated as follows: MPE*_product_*/MPE*_precursor_*.

#### Formation of IBD panel cell

We make the myeloid progenitor cell according to the protocol as described previously^53^. In summary, we collected bone marrow from the femur and tibia of female SAMP1/YitFc mice and carried out a negative selection process to isolate lineage-negative progenitor cells. This selection was accomplished using a cocktail of antibody that specifically targeted Thy1.2, B220 and MacI markers. Subsequently, we separated the lin+ cells from the progenitors using a Stemcell Technologies magnetic column. The isolated progenitor cells were then subjected to a 48-hour pre-stimulation in a culture medium consisting of Iscove’s Modified Dulbecco Medium (IMDM) with 15% fetal bovine serum (FBS), 1% penicillin-streptomycin-glutamine (PSE; Gibco), and supplemented with 50 ng/ml of stem cell factor (SCF), 25 ng/ml of interleukin-3 (IL-3), and 25 ng/ml of interleukin-6 (IL-6).For the introduction of the ER-Hoxb8 retrovirus, we infected 25,000 marrow progenitors with 1 ml of the retrovirus through a process known as spinoculation, which involved subjecting the cells to 2,500g of centrifugal force for 2 hours at 22°C. Lipofectamine was used in conjunction with the retrovirus at a ratio of 1:1,000 (Gibco), with detailed instructions provided in the Supplementary Methods. Following the infection, the progenitors were cultured in OptiMem medium, which included 10% FBS, 1% PSE, 10 ng/ml SCF, and 30 μM β-mercaptoethanol (1 μl per 500 ml of medium), along with 1 μM β-estradiol (estrogen; Sigma). Based on the initial rates of progenitor outgrowth in the presence or absence of G418 selection, we estimated an infection efficiency of approximately 10%. To enrich immortalized myeloid progenitors, we regularly transferred nonadherent cells into new 12-well tissue culture plates every 3 days. By day 14, these immortalized progenitors became the dominant cell type in the cultures, with a cell generation time of approximately 18-20 hours. In contrast, control cultures exhibited reduced proliferation and ceased dividing by day 21. It’s important to note that if GM-CSF was employed instead of SCF during the selection of immortalized progenitors in this protocol, it resulted in the emergence of clones capable of differentiating into both neutrophils and macrophages.

#### Cell viability

The SAMP myeloid progenitor stem cells were cultured in DMEM (Dulbecco’s Modified Eagle’s Medium (DMEM)) and supplemented with 10% FCS, penicillin–streptomycin, glutamax, and non-essential amino acids (NEAA). After six days, cells were plated for experiment. 1X10^4^ cells were plated on the 384 well plate. The *E. coli* and *Parabacteroides distasonis* were cultured and suspended in PBS. Each bacterial extract was heat killed for 30 min at 95°C. The OD600 value was obtained to normalize the concentrations of each bacterial suspension. The cells were treated with the 10 µl bacterial extract for 18 hours. The cell viability assay was performed by CellTiter-Glo® 3D (Promega). The assay was conducted according to the manufacturer’s protocol. In short, after 24 hours, the same volume of reagent was added to the cell, and they were incubated for 30 minutes. After that, luminescence reading was done by SpectraMax i3x.

#### Multiple sequence alignment for presenting the SNP

We picked the sequence on the basis of genome rearrangement as shown in **Supplementary Figure 2A** and we used these sequences in the blast query to find the aligned sequence of CavFT46. The multiple sequence alignment is done by the software CLC Sequence Viewer 8.0.0 under the default setting to find SNP, Gap and insertion.

#### Data Availability

The genome of the *P. distasonis* CavFT-hAR56 strain is available in Gene Bank accession for CavFT-hAR56 is <u>CP124739.1</u>, BioSample <u>SAMN11642307</u>, <u>SAMN30072245</u>; SRA <u>SRS14371963</u>. National Center of Biotechnology Information used Prokaryotic Genome Annotation Pipeline for the annotation of the genome. This pipeline uses HMM (hidden Markov models) for functional annotation of proteins (assembly name **<u>ASM2432872v1</u>**;www.ncbi.nlm.nih.gov/genome/annotationprok/).

## Acknowledgements

This project was supported by NIH grant R21 DK118373 to A.R.-P., entitled “Identification of pathogenic bacteria in Crohn’s disease.” We acknowledge the Biorepository Core of the NIH Silvio O. Conte Cleveland Digestive Disease Research Core Center (P30DK097948) for clinical advice and provision of deidentified surgical specimens for the above-mentioned R21 project. Partial support earlier originated for A.R.-P. from a career development award and recently from a Litwin Pioneers award from the Crohn’s and Colitis Foundation. The bowel specimens from which the present isolates were obtained for clinical research purposes were acquired as de-identified surgical specimens CD patients via the NIH Silvio O. Conte Cleveland Digestive Disease Research Core Center in Cleveland, Ohio, USA, in compliance with IRB-approved protocols (IRB protocol number: 02-10-07) authorized by the University Hospitals Cleveland Medical Center and Case Western Reserve University review boards. Special thanks to Dr. Weibiao Cao, Professor of Pathology and Laboratory Medicine, Brown University, RI, USA, for scientific criticism and suggestions on the integration of mucosal fissures and fistulas biology.

## Authors Contribution

ARP conceptualized and initiated all studies. VS and ARP conceptualized and conducted the genomic analysis and drafted the manuscript. GW, KT, JF, ARB and FC assisted with human sample collection and established IRB protocols for the Cleveland Digestive Diseases Research Core Center Biorepository to enable the study. CF, SEV and BE supported the MRI experiments. IR oversaw and conducted the target metabolomics experiment. NCB, ARKD, XY, and AV contributed to genome analysis. CEG and MRJ conducted the bacteria identification experiments. BG, MW, PM and SA were involved in the mouse experiment, while DS, CM, JE and MS participated in the cell experiment ARP leads the program for identification of pathogenic bacteria in severe surgical Inflammatory Bowel Disease/Crohn’s Disease. All authors approved the final version of the manuscript.

