## Supplementary Data for "Clonal *Parabacteroides* from Gut Microfistulous Tracts as Transmissible Cytotoxic Succinate-Commensal Model of Crohn’s Disease Complications"

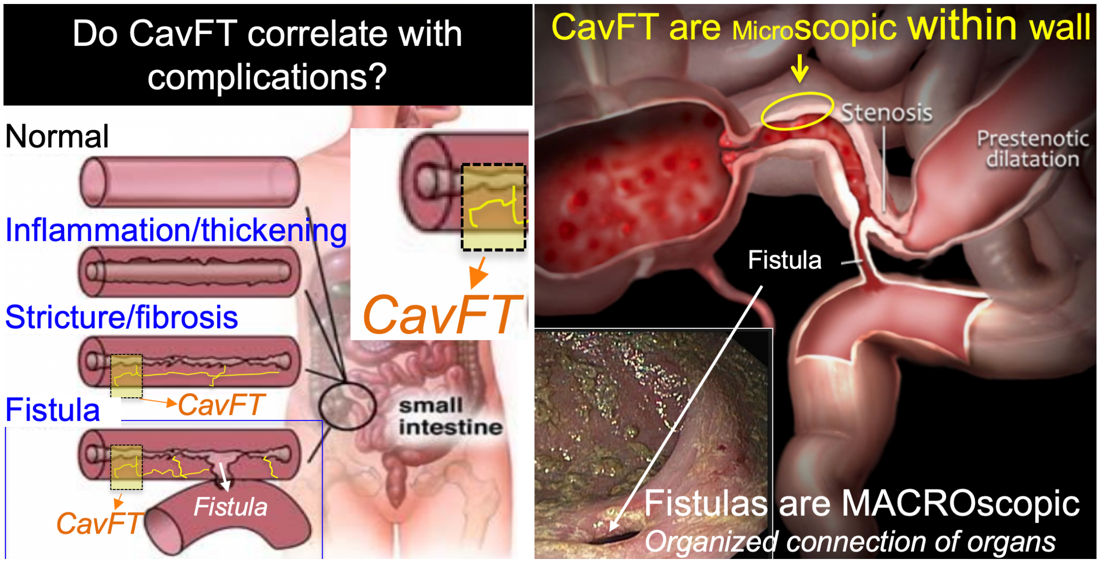


**Supplementary *Figure 1:*** How CavFT could be the etiological & mechanical precedent to fistulas.

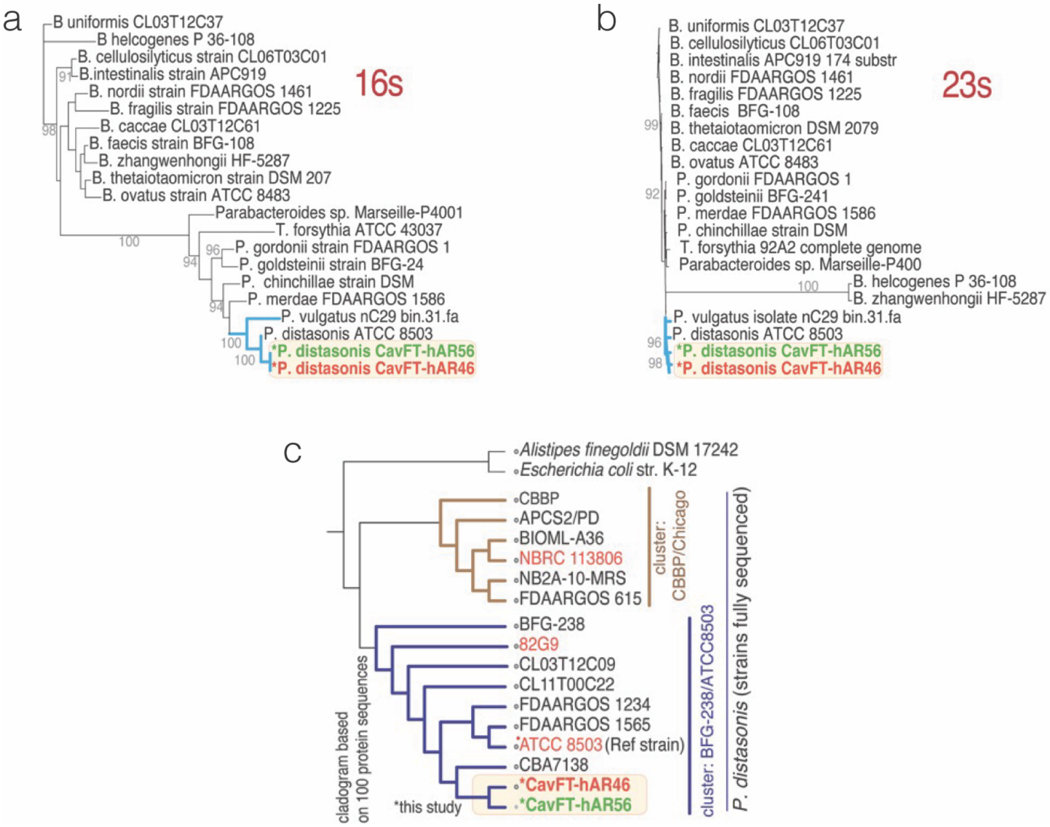


**Supplementary Figure 2: *P. distasonis* CavFT strains cluster together:** Maximum likelihood phylogenetic analysis of **a)** 16s, b) 23s ribosomal proteins and **c)** Phylogenomics of *P. distasonis* protein-coding sequences. **Supplementary Table 1,** genome sequencing statistics for both CavFT isolates. Note protein CavFT supercluster. CBA7138 (South Korea, healthy human feces, 2018). One called the '**CBBP/Chicago-protein-cluster'**, and the second, **'CL11T00C22/ATCC8503-protein-cluster'** (CL11T00C22, human-feces, USA, 2009), which harbors the CavFTs as a clade with strain CBA7138.

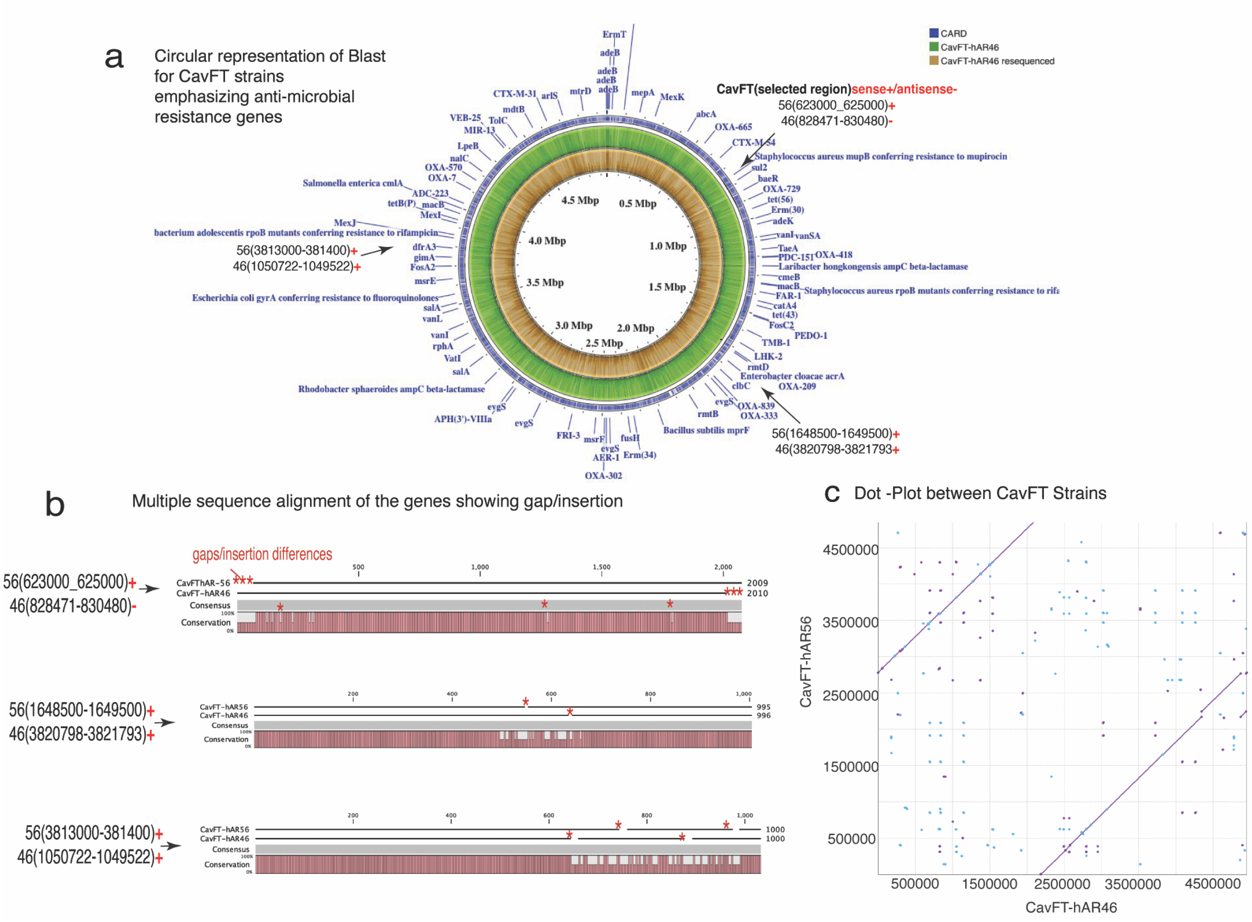

**Supplementary Figure 3.** ***P. distasonis*** **CavFT strain are different with each other** **a**) Circular representation of draft genome and features of CavFT-hAR56 using Prokka Annotation and we blast the CavFT-hAR46 (published and re-sequenced) with CavFT-hAR56. The figure were produced using CgView. **b)** MSA of on the basis of blast results showing insertion and deletion in the alignment and **c)** Heat map showing antimicrobial susceptibility of CavFT strain with *Pediococcous acidilactici* and *Enterococcus faecium*. We used the Morpheus broad institute for formation heat map(https://software.broadinstitute.org/morpheus/) **c)** Dot plot between *P. distasonis* CavFT-hAR46 and CavFT-hAR56

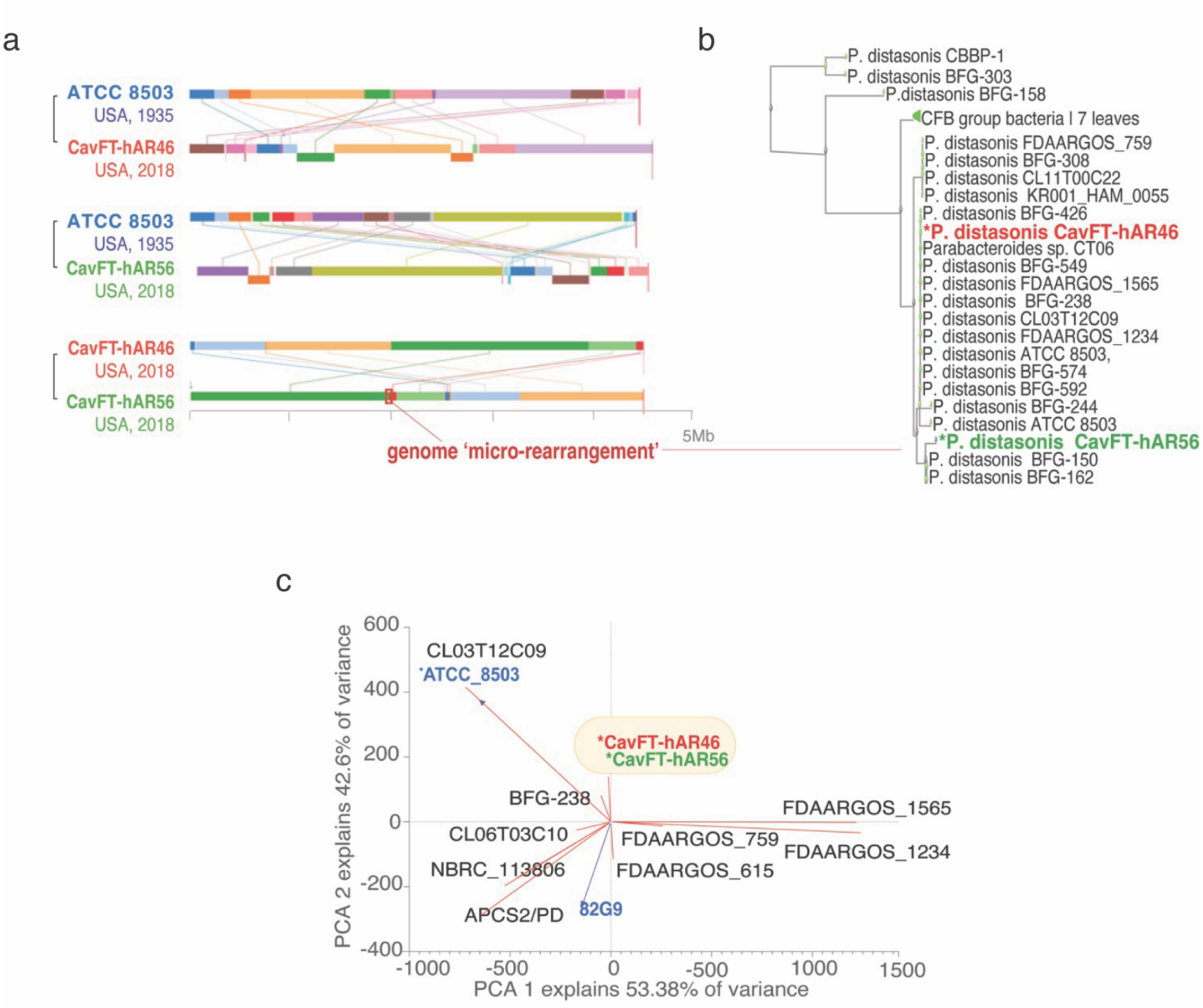


**Supplementary Figure 4. Genome rearrangement show difference in CavFT Strains. a)**Mauve genome pairwise alignment of *P. distasonis* ATCC 8503 (1935), CavFT-hAR46 and -hAR56 (2017). Distinct colors illustrate rearranged genome areas. CavFT-pairwise shows few rearrangements, indicating strains have evolved separately. **b)** Selected CavFT micro-rearrangement sequence used for phylogenetics shows CavFT strain differences. **c)** PCA and biplot loadings of genome rearrangement distances within complete genomes of *P. distasonis* vs. ATCC 8503 and 82G9 using as reference strains. CavFTs distinct similar rearrangements.

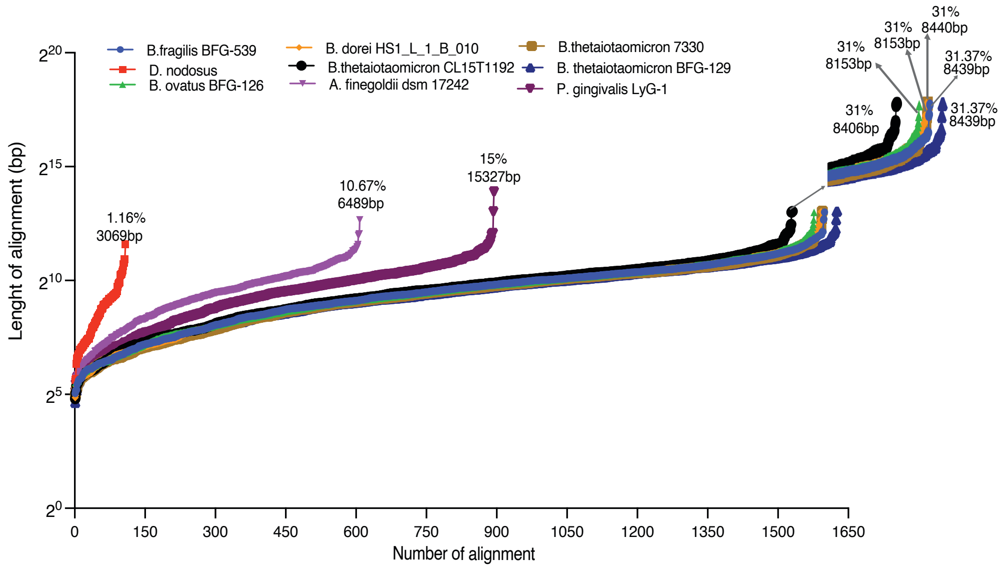


**Supplementary Figure 5.** 'Claw plot' illustrating the ranking of DNA fragments that ATCC8503 shares with other genomes. *Bacteroides* spp.

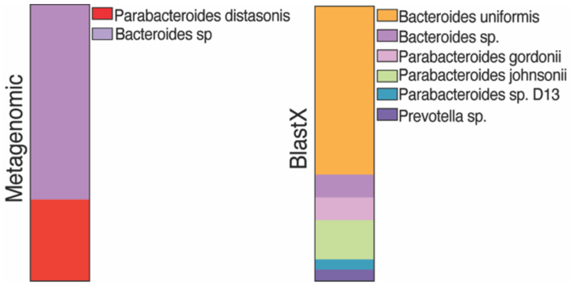


**Supplementary Figure 6.** Metagenomics and BlastX analysis of the 56575bp CavFT/*Bacteroides* insert CavFT-hAR56 split into 250bp fragments illustrates numerous *Bacteroides* spp. GO terms**.**

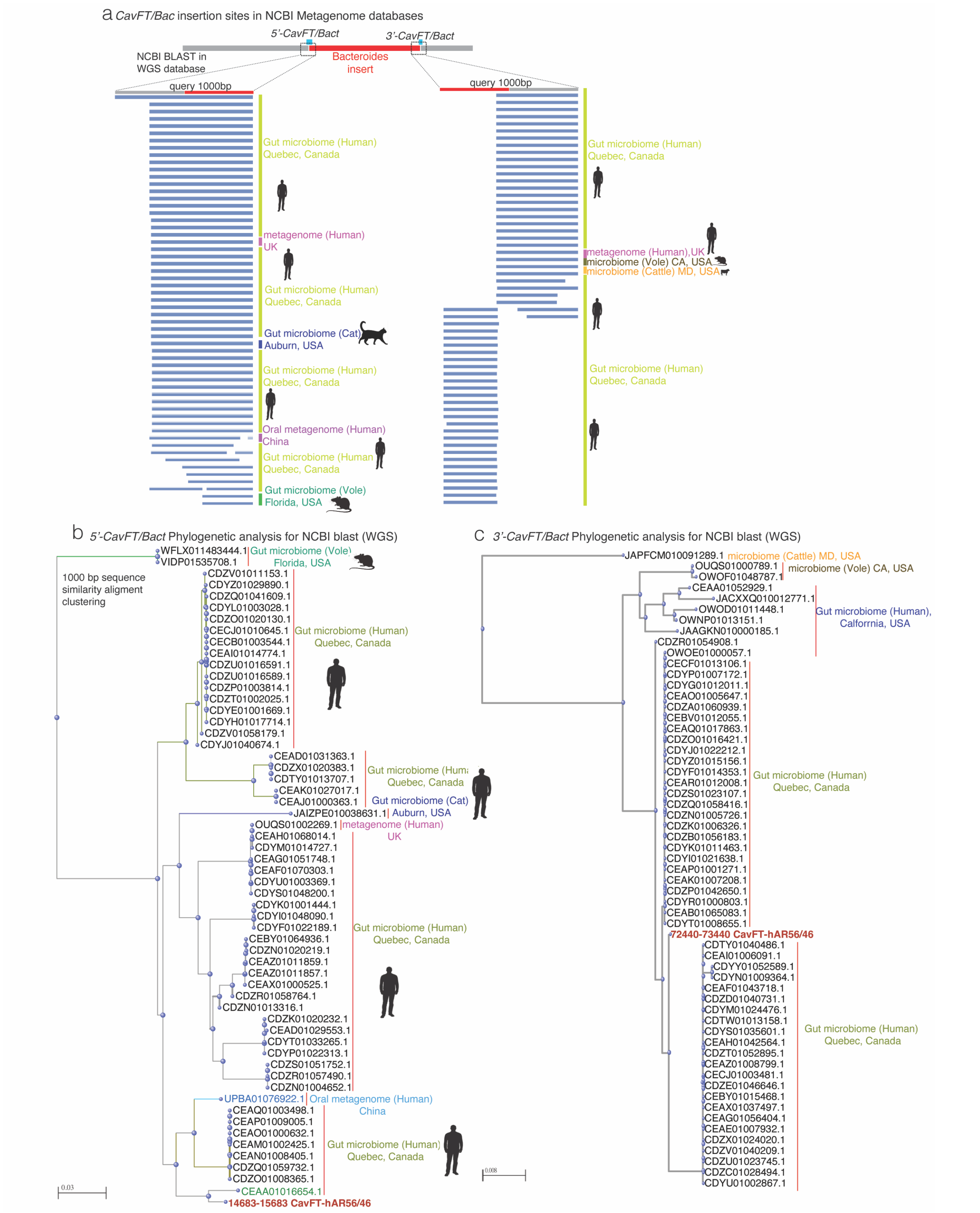


**Supplementary Figure 7.: a)**The Multiple Sequence Alignment (MSA) of the BLAST hit was conducted using Whole Genome Sequencing (WGS) data available on NCBI. We focused on the 1000 bp regions from both 5'-CavFT/bac and 3'-CavFT/bac. NCBI-Blast Phylogenetic tree for 1000 bp regions from both **b)**5'-CavFT/bac **c)** 3'-CavFT/bac

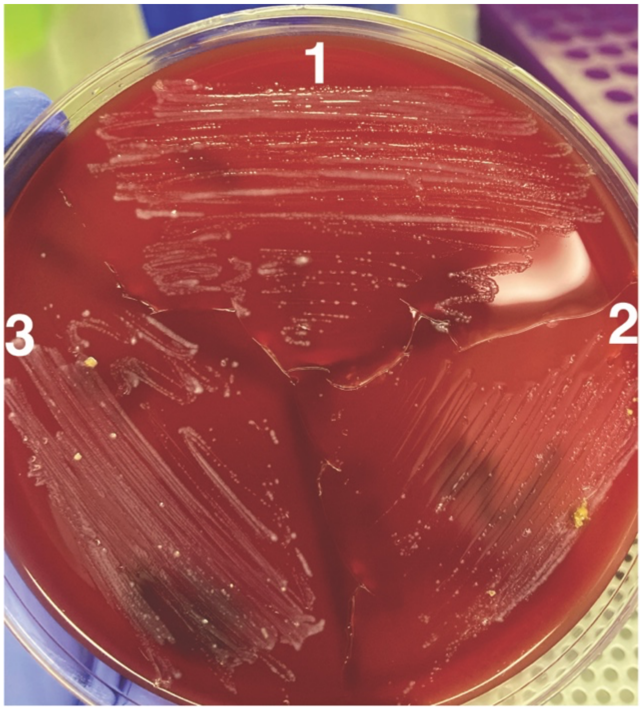


**Supplementary Figure 8.** We administered the bacteria through gavage to GF-SAMP mice 1, and the results indicate that GF-SAMP mice 2 and 3 were also contaminated by the first mouse, as observed in the plates.

**Supplementary Table 1:** The genomic characteristics of *Parabacteroides distasonis* strains CavFT-hAR46 (re-sequenced) and CavFT-hAR56 were compared to those of other complete *P. distasonis* genomes

| Feature | Parabacteroides distasonis Strain | | | | | |
|---|---|---|---|---|---|---|
| | **CavFT56*** | **CavFT-hAR46 [reproducibility]** | **ATCC 8503** | **FDAARGOS _759** | **NBRC 113806** | **82G9** |
| Origin (year) | Ohio, USA | Ohio, USA (2019) | USA | USA | - | Japan |
| Accession no. | CP124739 | CP040468 | NC_009615 | NZ_CP054012.1 | NZ_AP019729 | NZ_LR215978.1 |
| Genome size (bp) | 4951235 | 4952323 [99.9994%] | 4811379 | 4926033 | 5179960 | 5212259 |
| No. of CDS | 4245 | 4263 [99.929%] | 4216 | 3980 | 4171 | 4365 |
| G+C content (%) | 45.2 | 45.2 [100%] | 45.1 | 45.2 | 45.1 | 45.2 |
| No. of rRNAs | 21 | 21 [100%] | 21 | 21 | 21 | 21 |
| No. of proteins with functional assignments | 2674 | 2694 [99.4%] | 2639 | 3403 | 2494 | ND |
| No. of hypothetical proteins | 1571 | 1569 [100%] | 1487 | 577 | 1677 | ND |
| No. of antibiotic resistance genes | 23 | 24 [100%] | 22 | 21 | 26 | ND |
| **CavFT-hAR56 compared to others** | | | | | | |
| Linear Coverage (%) | | 99.97 | 84.92 | 84 | 77.97 | 79.88 |
| Average Identity (%) | | 95.82 | 96.17 | 96.07 | 95.72 | 96.14 |

**Supplementary Table 2**. Transmural lesion for of intestinal specimens: terminology.[1]

| |
|---|
| **SM-cobblestone (SmCobbles).** Aggregated villi in the small intestine or bulking lesions in the colon, corresponding to the equivalent cobblestone lesions observed during endoscopy in humans, can be also referred to as cobblestones in stereomicroscopy. |
| **Mucosal fissures (SmFissures).** Lesions that correspond to cracks in the epithelium accompanied with areas of abnormal discoloration or texture in the mucosa/submucosa areas that correspond to the confluence of two cobblestone lesion curvatures |
| **SM congestion lesions (SmCL).** SmCL are areas with high red discoloration and visible blood vessels compared to surrounding tissue, similar to SmLL (see below) discoloration when tissues are moistened with 70% ethanol, but that appear dry upon evaporation of the ethanol. SmCL may represent areas of early inflammation with predominant vascular congestion (and no tissue liquefaction as SmLL). |
| **SM liquefaction lesions (SmLL).** Resemble areas of 'melting' (liquefaction) of tissues whose solid structure disappears due to inflammation or tissue degradation. Areas have red discoloration and contain more visible blood vessels. Upon dehydration of formalin-fixed ethanol-preserved tissue, SmLL lesions maintain a moistened appearance with respect to their surroundings, suggesting physico-chemical tissue property changes resulting from inflammation. SmLL represent areas where severe infiltration of immune cells has replaced matrix structure and matrix forming cells. They contain higher cellularity (host nuclear DNA copies) if determined by B-actin qPCR CT values on DNA extracts (not RNA/cDNA). SmLL affect any gut tissue layer. |
| **Penetrating fistulous tracts (PFT).** Linear irregular-edge lesions that resemble narrow tracts running perpendicular to the gut lumen alongside muscle bundles of the circular muscle layer. They run from (sub)mucosal layers to reach the deep longitudinal muscle layer. The PFT may represent the earliest form of dissecting inflammatory conditions that communicate (sub)mucosal layers with deeper structures such as muscular or serosa layers. |
